# Biomimetic Artificial Bone Marrow Niches for the Scale Up of Hematopoietic Stem Cells

**DOI:** 10.1101/2023.06.23.546233

**Authors:** Minerva Bosch-Fortea, Daniele Marciano, Julien E. Gautrot

## Abstract

Hematopoietic stem cell (HSC) transplantation to treat haematological disorders is greatly restricted by poor cell availability. *Ex vivo* expansion of HSCs as a strategy to overcome this limitation has shown limited success so far, due to the loss of stem cell properties in culture. Therefore, engineering of culture platforms that mimic the physiological properties of the bone marrow (BM) in a scalable format is an important target to enable the translation of HSC therapies. Here, we report the design of biomimetic BM niches that enable the culture of HSCs in a scalable 3D platform. Beyond cellular and biochemical components (e.g., matrix and growth factors), an important element of the BM microenvironment is its architecture, dense in adipocytes, with relatively limited matrix and anisotropic mechanical properties. To capture this context, we propose the use of bioemulsions^1,2^ in which oil microdroplets and associated mechanical anisotropy recreate important architectural features of the hematopoietic niche. Mesenchymal stem cells (MSCs) grown at the surface of such bioemulsion remodelled this environment, assembling an interstitial matrix mimicking that of the BM microenvironment composition. In addition, MSCs secreted important factors underpinning the crosstalk between stromal cells and HSCs in the native environment. HSCs cultured in the resulting artificial BM niches maintained stemness whilst expanding significantly (> 33-fold compared to HSCs cultured in suspension) and enabling scale up of expansion in conical flask bioreactors, to produce 2M cells in a single batch. This platform therefore harnesses engineered BM microenvironments and the processability of bioemulsions and microdroplet technologies to produce HSCs in a scalable format, for application in cell-based therapies.

## 1. Introduction

Hematopoietic stem and progenitor cells (HSPCs), which mostly reside in the bone marrow (BM), give rise to all blood cell lineages and are capable of self-renewing, underpinning the homeostasis of haematopoiesis. This capacity is directly targeted by cell therapies aiming to treat haematological disorders (such as various types of cancers and autoimmune diseases), as HSPC transplantation allows the reconstitution of a healthy blood population^1,2^. However, the clinical use of HSPCs has been hindered by both the scarcity of compatible donors and the low concentration of cells in donor tissues^3^, with mobilised peripheral blood (mPB) currently being the main source for HSPCs^4^. Since the availability of a larger cell population could improve clinical outcomes^5^, the *ex vivo* expansion of HSPCs has been identified as the most promising approach to overcome suboptimal stem cell dose during transplantation^6^. This would not only improve the outcome of autologous and allogenic transplantation of mPB HSPCs but will potentially expand the use of HSPCs isolated from cord blood (CB) to adult patients. However, enhancing the availability of transplantable HSPCs prior to infusion was found to be particularly challenging because the complexity of the hematopoietic niche makes it difficult to unpick the intricate interactions regulating HSPC biology^7^. The BM niche is a heterogenous complex microenvironment that includes cellular (stromal cells), biochemical (growth factors and cytokines), and physical (elastic and shear moduli, viscoelasticity, microstructure, oxygen tension) factors regulating HSPC behaviour^8,9^. The dynamic orchestration of the balance between these different factors is thought to underpin the regulation of HSPC function (i.e., quiescence, proliferation, migration, differentiation and apoptosis), and is difficult to capture *in vitro*.

Significant efforts have focused on defining and improving appropriate culture conditions to stimulate *ex vivo* expansion of ^6,10^. Critical to this issue is the maintenance of stem cell properties as HSPCs undergo rapid differentiation in culture, leading to the loss of their reconstituting and therapeutic potential^11^. Likewise, the discrepancy between HSPC behaviour *in vitro* and in physiological conditions makes it particularly challenging to gain deeper understanding of HSPC biology and the numerous processes potentially missregulated during malignant transformation of the BM niche^12^. These pitfalls have prompted the design of engineered microenvironments able to mimic some of the properties of the hematopoietic niche, to reproduce HSPCs phenotype *in vitro*^13–15^. A number of procedures have proposed to mimic cellular interactions of HSPCs with adjuvant stromal cells through co-culture with BM-derived mesenchymal stem cells (MSCs)^16–19^, endothelial cells^20^, and other stromal cell types^21^. Others aimed at modelling the acellular microenvironment through the engineering of biochemical and biophysical components of the niche^22,23^. However these various platforms remain difficult to scale-up, often lack efficiency or are difficult to process to effectively extract HSPCs for reimplantation^24^, especially considering the scale required for proper engraftment, typically requiring doses of 2 to 8 x 10^6^ cells/kg CD34^+^ ^25^.

Here we report the design of artificial BM niches (@BMNs) that recapitulate the microstructure, mechanical anisotropy and some of the biochemical and cellular complexity of the BM HSPC microenvironment. This platform is based on bioemulsions, in which protein nanosheets stabilise microdroplets and confer strong and elastic interfacial shear mechanics^26,27^ mimicking the anisotropy of the interstitial extra cellular matrix lining adipocytes in the BM. Bioemulsions enable the rapid expansion of MSCs, which prime the formation of @BMN through significant matrix remodelling to form microtissues and the acquisition of a pro-hematopoietic secretory phenotype. In turn, these niches promote the expansion of HSPCs whilst maintaining their phenotype and capacity to commit to multiple hematopoietic lineages. Finally, the scale up (100-fold) of this platform to conical flask bioreactors typically used for the culture of planktonic cells and cell-seeded microcarriers is presented, with excellent retention of HSPC expansion (> 33-fold compared to suspension culture of HSPCs) and phenotype.

## 2. Results and Discussion

### Design of a biomimetic artificial bone marrow microenvironment

The design of the @BMN aimed to capture several key features of the BM microenvironment. Among them, the architecture of the BM is particularly important for the regulation of HSPC behaviour. The BM niche mainly consists of collagen and fibronectin networks heterogeneously distributed and wrapped around adipocytes (the most abundant cell type in the BM), providing an overall soft environment (E = 0.25-24.7 kPa, although reports vary considerably)^28^. Although adipocytes have recently been reported to contribute to hematopoiesis through secretion of SCF and other cytokines and adipokynes ^29^, they are often not considered as critical signalling hubs regulating HSPC phenotype^30^. However, adipocytes have a strong impact on the structure and mechanical anisotropy and heterogeneity of the BM environment^31,32^. From a structural and mechanical point of view, adipocytes may be regarded as oil microdroplets surrounded by a viscoelastic membrane composed of the cortex, lipid bilayer and pericellular matrix^33,34^. MSCs find themselves nested within this microenvironment, contributing to its remodelling and providing additional factors and secretion for the maintenance a healthy HSPC niche^35^ (Figure 1A). Therefore, we proposed to engineer @BM based on protein nanosheet-stabilised microdroplets that capture some of the microstructural properties and mechanical anisotropy of the BM niche (Figures 1A and B). Indeed, protein nanosheets were recently found to support the adhesion of adherent cells, including MSCs, and the maintenance of the phenotype, supporting the long-term expansion of MSCs and the retention of their capacity to differentiated in defined lineages^36,37^. Such processes were found to be associated with the strong viscoelastic properties of self-assembled protein nanosheets, and their ability to resist cell-mediated contractile forces^26,27^. Such assemblies are inherently associated with high mechanical anisotropy, with high interfacial shear moduli (in the plane of the interface) and low transverse moduli^27,38^, therefore capturing some of the features of the BM microenvironment.

**Figure 1.**
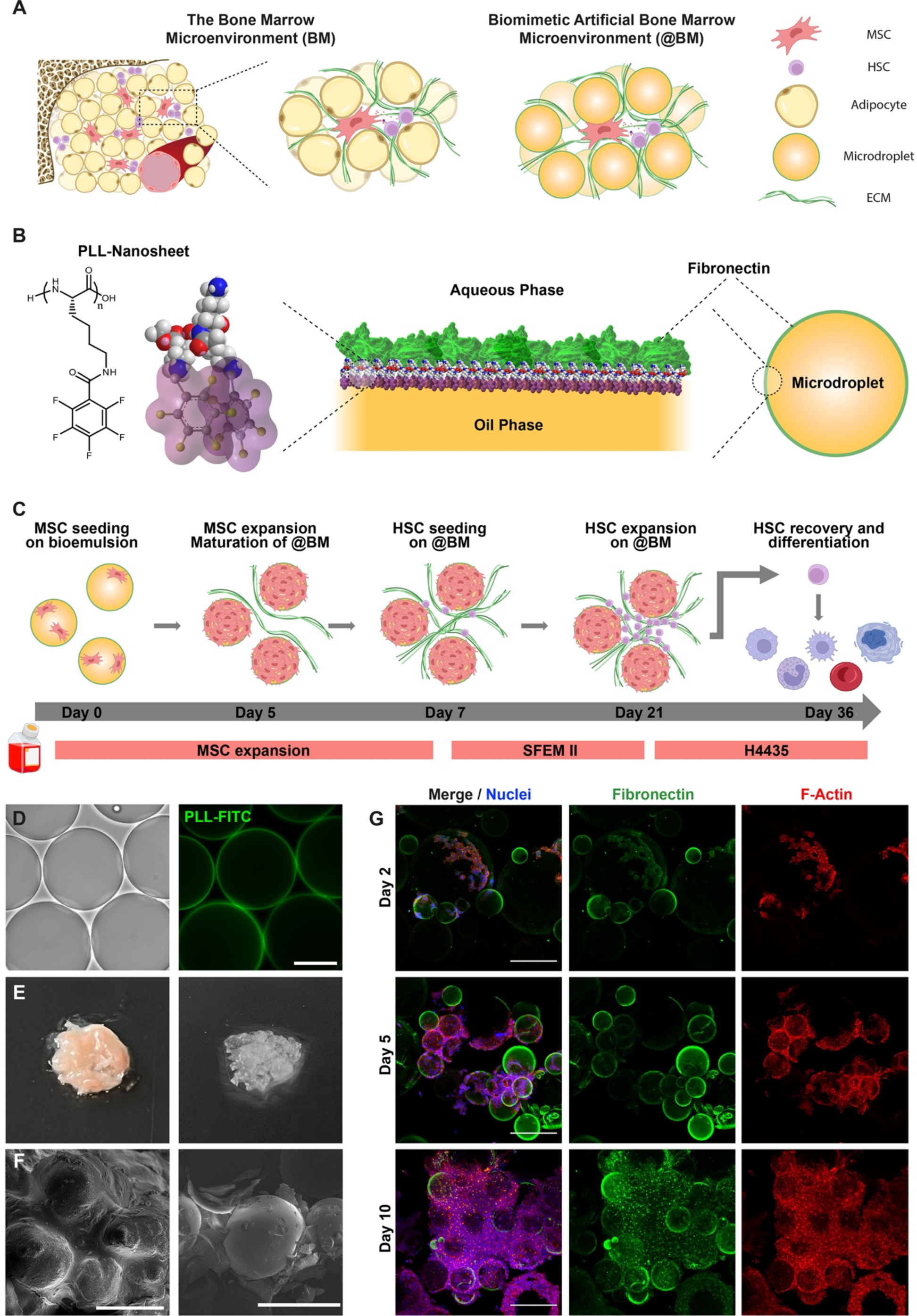
Design of a biomimetic artificial BM microenvironment. A) Schematic representation of the BM microenvironment and the proposed architecture and composition of @BMN for *in vitro* HSPC culture. B) Schematic representation of poly(L-lysine) nanosheets, to stabilise microdroplets and enable fibronectin adsorption. C) Detail of the protocol used to assemble @BMs. MSCs are seeded at day 0 and cultured in MSC growth medium for 7 days to allow cell expansion and maturation of a BM biomimetic niche. On day 6, the growth medium is switched to SFEM II medium supplemented with 100 μg/mL SCF and on day 7, HSPCs are added to the culture and further expanded for 14 days. On day 21, CD34^+^ cells are isolated from the @BM and their phenotype and differentiation potential assessed. D) Images of oil droplets (left, brightfield) and PLL nanosheets assembled at corresponding interfaces (Alexa Fluor 488-tagged; right). Scale bar, 50 μm. (E) Images of bovine BM (left) and @BM after 15 days of MSCs culture (right). (F) Cryo-scanning electron micrographs from bovine BM samples (left) and bioemulsion prior to cell seeding (right). Scale bars, 300 μm (left) and 500 μm (right). (G) z-projections of confocal z-stack of developing @BMs, with MSCs growing on emulsions using MSC medium for 2, 5 and 10 days. Scale bar, 200 μm.

Bioemulsions stabilised by poly(L-lysine) (PLL) nanosheets and enabling the capture of fibronectin were used as basal system for the conditioning of @BMN (Figures 1B and C). The homogenous coating of microdroplets by PLL nanosheets was confirmed by fluorescence microscopy (Figure 1D). The resulting emulsion, with mean droplet diameters of 155 ± 6 μm, displayed excellent stability over a period of 15 days upon incubation, with modest increase to 196 ± 95 μm (Supplementary Figure S1). MSCs were seeded on bioemulsions and allowed to proliferate for 15 days, under gentle agitation, during which time they gradually remodelled this environment and aggregated individual microdroplets into microtissues (Figure E). The microstructure of these assemblies was characterised by cryo-SEM and was found to resemble that of bovine BM, with apparent adipocyte-mimicking oil microdroplets separated by interstitial matrix (Figure 1F). Fluorescence microscopy confirmed the gradual proliferation of MSCs at the surface of microdroplets and the aggregation of multiple droplets to form microtissues (Figure 1G). Whereas initially (days 1-5) MSCs adhered and proliferated at the surface of the droplets, over prolonged culture times they formed interstitial tissues that connected multiple droplets (Figure 1G and Supplementary Figure S2). After 5 days in cultures, MSCs were found to grow both at the surface and in between droplets and the prevalence of this interstitial tissue was clearer after 10 days of culture.

### Preservation of the MSC phenotype during the shaping of artificial BM microenvironments

The key role that MSCs play in the native hematopoietic niche^18,39^ positions MSC/HSPC co-cultures as a valuable approach to mimic the physiology of the BM. Thus, we explored the maintenance of MSCs phenotypic traits during the development of @BMN. Considering the importance of integrin-mediated adhesion to MSC phenotype, we first investigated MSC spreading at the surface of bioemulsions (Figure 2A). Similar to MSCs spreading on glass or other rigid substrates (Supplementary Figure S3), MSCs assembled a structured F-actin cytoskeleton at nanosheet-stabilised liquid-liquid interfaces. In addition, these assemblies were also associated with the formation of mature focal adhesions, characterised by β1-integrin clustering and vinculin recruitment, at the end of apparent stress fibres (Figure 2A). This is consistent with previous observations that incubation with a β1-integrins function-blocking antibody inhibits the spreading of MSCs and that their spreading at nanosheet-stabilised liquid interfaces is affected similarly by disruptors of actin assembly (e.g., blebbistatin and ROCK inhibitor) to cells spreading on rigid substrates^37,40^.

**Figure 2.**
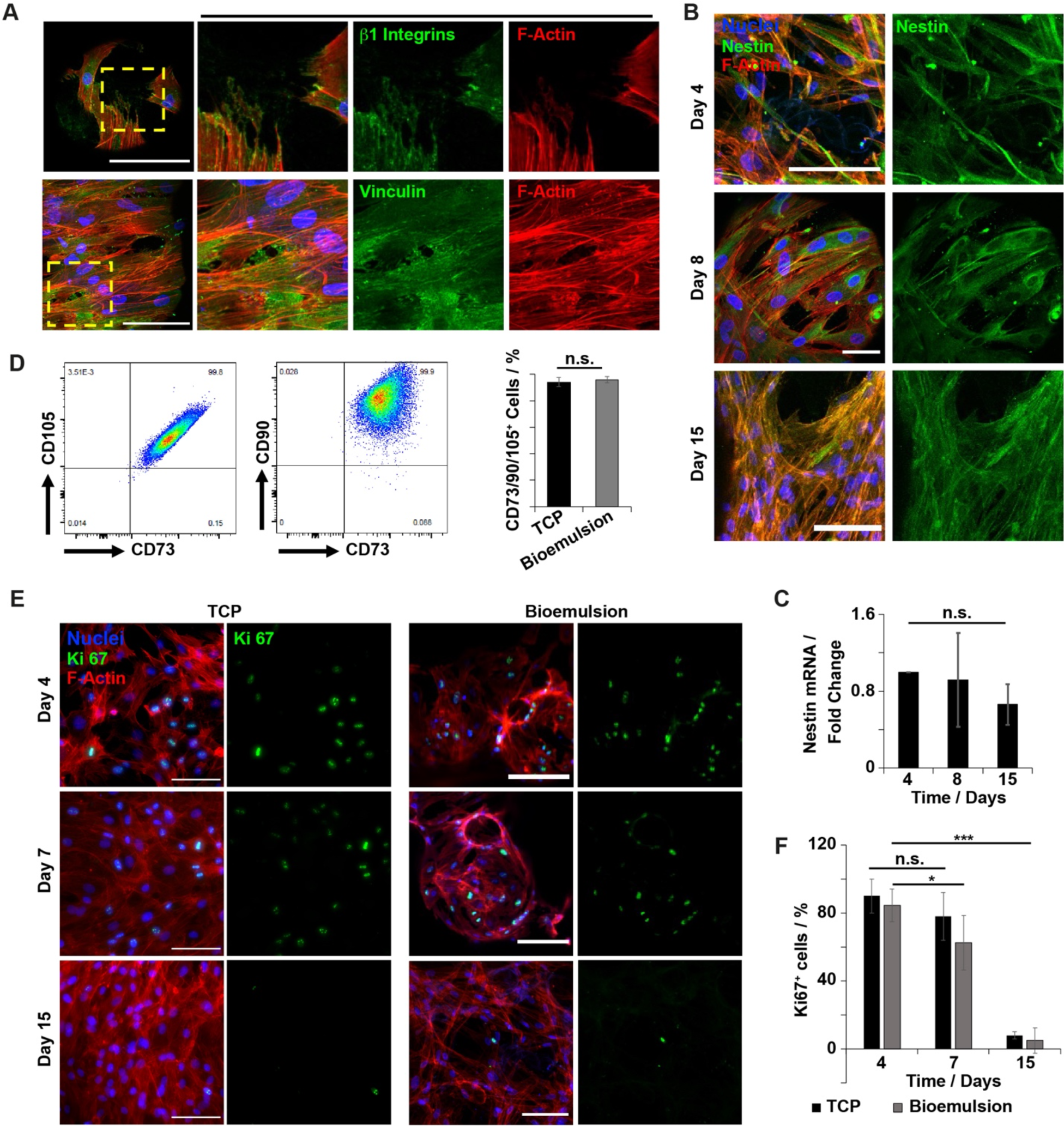
The stemness of MSCs is maintained in the developing @BMN. A) Confocal images of MSCs adhering to PLL nanosheet-stabilised bioemulsions, demonstrating the formation of mature focal adhesions (ß1-integrin and vinculin, green) and stress fibres (F-actin, red) after 2 and 15 days respectively in culture on bioemulsions. Scale bar, 20 μm (upper), 50 μm (lower). (B) z-Projections of confocal z-stacks of MSCs growing on bioemulsions at different time points (in MSC medium), confirming the maintenance of Nestin expression (green). Scale bar, 50 μm, 20 μm and 100 μm respectively. C) Quantification of Nestin expression at the mRNA level, relative to levels at day 4. Mean ± SD; N = 3; n.s., P > 0.05. (D) Gating strategy (left) and percentages of cells displaying triple marker expression (CD73, CD105 and CD90; right), comparing MSCs growing on 2D tissue culture plastic (TCP) and on bioemulsions in MSC medium, at day 15. Mean ± SD; N = 3; n.s., P > 0.05. After doublet exclusion, mononuclear cells were gated based on low/high expression of calcein to identify live cells and exclude bioemulsions and dead cells. CD73^+^/CD90^+^ subset was selected and CD105^+^ cells were subsequently gated to identify the population of interest. (E) z-Projections of confocal z-stacks of Ki-67 immunostained samples of MSCs growing on 2D substrates (left) or on bioemulsions (right), at different time-points in MSC medium. Scale bars, 50 μm, (F) Corresponding quantification of percentages of Ki-67^+^ MSCs. Mean ± SD; N = 3; n.s., P>0.05; *, P < 0.05; ***, P < 0.001).

To further establish how culture at liquid-liquid interfaces preserved MSC phenotype, we explored the expression of Nestin, an important stemness marker associated with the maintenance and homing of HSPCs in the BM niche^41^. Analysis of Nestin expression at different time-points confirmed its maintenance over 15 days of culture (Figures 2B and C). Flow cytometry further established the preservation on bioemulsions of triple marker (95.8 ± 2.4% CD105^+^CD73^+^CD90^+^ population) expression throughout the priming and early stage of HSPC culture, within the niche (15 days, Figure 2D). These observations are in good agreement with the maintenance of MSC stemness during the long-term expansion of MSCs on bioemulsions^36^. In addition, Ki-67 immunostaining indicated a reduction of cycling, in particular after 7 days of culture, as MSC confluency inhibited proliferation and triggered cycling arrest (Figures 2E and F)^42^. This restriction in cell proliferation will ensure the maintenance of MSCs cell pool to low levels on the long-term without the need of further treatment (e.g., irradiation) while it would not impact the proliferation of HSPCs since these cells grow at the surface of MSCs and in suspension.

### MSCs remodel and prime artificial BM niches

Several studies have highlighted the crucial role of the ECM in signalling and homing mechanisms occurring in the hematopoietic niche. This regulation is proposed to take place through the direct engagement of ECM components by specific surface receptors in HSPCs^43,44^ or through sequestration/release of specific growth factors, morphogens, and cytokines^45,46^. Such matrix signalling was particularly evidenced by the higher expansion levels observed when HSPCs were grown on acellular matrices derived from BM stromal cells^47^. Therefore, we next investigated whether MSCs assembled a BM-like ECM upon culture on bioemulsions. Cryo-SEM of bioemulsions freshly generated and after culture of MSCs for 15 days (but following cell detachment) gave evidence for the formation of matrix fibre deposition at the liquid-liquid interface. Indeed, whereas relatively amorphous featureless nanosheets could be observed on pristine microdroplets, in agreement with prior imaging via TEM and SEM^26,27^, bundles of nanofibers could be clearly observed at the surface of droplets following 15 days of culture (Figure 3A). This phenomenon was further confirmed by immunostaining, revealing the formation of dense mats of fibres of fibronectin and laminin (Figure 3B). Unlike the fibronectin initially adsorbed at the surface of PLL nanosheets, the fibronectin matrix observed at later time points displayed a clear nanofibrous morphology, typical of physiological fibronectin-rich matrices. This matrix was also observed in areas not occupied by cells (Supplementary Figure S4), therefore implying that it is not simply associated with pericellular matrix but contributes to the formation of an integrated microenvironment at the micro-to-mesoscale. The resulting ECM networks may stem from the remodelling of the fibronectin initially adsorbed at the surface of nanosheets or may result from *de novo* deposition of matrix. However, collagen I was found to be expressed but not to contribute to fibrillar networks within @BMN. Fibronectin is ubiquitously expressed across the BM and is required for long-term haematopoiesis and proliferation of HSPCs^48,49^, whereas laminin is involved in the homing and cell cycle regulation of HSPCs^50^. In contrast, although collagen I supports the growth of HSPCs, it is mainly found in the endosteal niche^51^. Therefore, key matrix proteins contributing to HPSC maintenance in the BM niche are integrated in the development of microdroplet-based @BMN.

**Figure 3.**
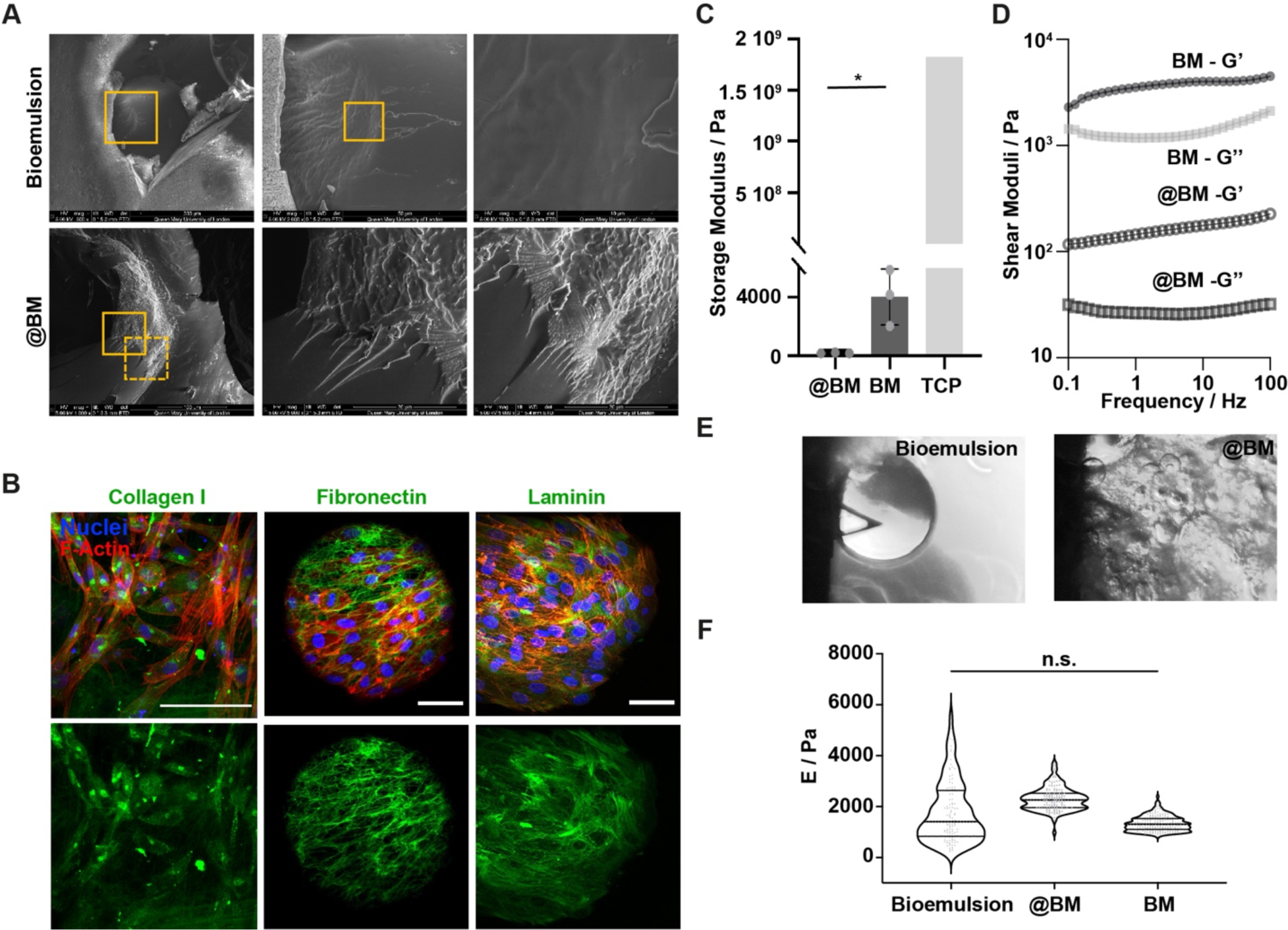
Matrix remodelling conditions the @BMN. A) Cryo-scanning electron micrographs of PLL nanosheet-stabilised bioemulsions prior to cell seeding (top) and after the culture of MSCs for 7 days (bottom). Yellow squares indicate areas magnified. Scale bar, 100 μm. (B) z-Projections of confocal z-stacks of MSCs cultured on bioemulsion microdroplets, immunostained to visualize ECM proteins. Scale bars, 50, 20 and 100 μm, left to right respectively. (C) Comparison of the shear storage modulus of @BM (day 14) and bovine BM measured by rheology, compared to TCP^55^ Mean ± SD; N = 3; *, P < 0.05. (D) Corresponding frequency sweeps recorded at a strain of 0.1%. (E) Images areas selected for nanoindentation of a bioemulsion interface and @BM sample. (F) Young’s moduli of nanosheet-stabilised bioemulsion interfaces, @BM and bovine BM tissue quantified via AFM indentation. Mean ± SD; N = 4-6; n.s., P > 0.05.

Associated with the assembly and remodelling of ECM networks, significant changes in matrix mechanics were anticipated. Topographical and mechanical cues are known factors regulating HSPC function and expansion^52–54^. A change in mechanics during the maturation of the @BMN is immediately evident from the rheological properties of bioemulsions. Whereas bioemulsions are fluid at early time points (preventing their characterisation via oscillatory rheology), MSC culture for 15 days led to the formation of microtissues that displayed more cohesive properties. At this time point, oscillatory rheology indicated a shear storage modulus of 180+25 Pa (Figures 3C and D). Whereas this is lower than the shear storage modulus measured for bovine BM (4000 ± 1600 Pa), such difference may partly be accounted for the difference in the dimension of @BMN, with microtissues collected typically spanning 0.5-1 mm, whereas bovine BM tissue of sufficient size to cover the 20 mm of our geometry could be obtained. Therefore, multiple @BMN had to be collated to carry out oscillatory rheology, potentially leading to long range relaxation and reducing the apparent moduli measured. In any case, the shear moduli measured for native and artificial BM were found to be relatively comparable and orders of magnitude lower than the Young’s modulus typically associated with tissue culture polystyrene (near 1.8 GPa ^55^).

Recognising the limitation of macroscale shear rheology to characterise the mechanics of microtissues, the local (nano-to-microscale) mechanical properties of BM microenvironments were examined using force probe microscopy (Figure 3E and Supplementary Figure S5). Nanoindentation of artificial and native BM tissues indicated comparable mechanical properties, with extracted Young’s moduli of 2300 and 2100 Pa, respectively (Figure 3F). However, it should be noted that Atomic Force Microscopy (AFM) alone does not enable the characterisation of the anisotropic mechanical properties of materials and that bioemulsions and native BM are expected to display significant anisotropy. The heterogeneity of the measurements is evident from the spread of the data gathered, compared to more homogenous substrates such as polyacrylamide hydrogels^56^. However, more strikingly, protein nanosheet-stabilised emulsions and microdroplets were recently reported to display orders of magnitude stiffer interfacial shear moduli than the transverse moduli associated with AFM indentation^38^. This is in contrast with surfactant-stabilised droplets, for which surface tensions, electrostatic interactions and van der Waals forces are reported to control the disjoining pressure of associated interfaces and AFM indentation profile^38^. Although BM constitute a more complex tissue than nanosheet-stabilised liquid-liquid interfaces, the nature of the matrix-adipocyte interface bears significant structural and mechanical analogy. Overall, our results indicate that the local mechanics and composition of key proteins of the BM niche are captured in the properties of @BMNs.

Additional important contributors to the supportive role of MSCs for the growth of HSPCs are their secretory phenotype as well as their ability to engage ligand-receptor interactions. For example, VCAM-1/VLA-4 interactions^57,58^ or interactions between CXCL12 and CXCR4^59,60^ are key to promote HSPC homing in the niche and contribute to maintain the HSPC pool. Indeed, inhibitors of both VLA-4 and CXCR4 are being used alone or in combination with G-SGF to mobilise HSPCs from PB in donors before transplantation^4,61^. Similarly, the production of Stem Cell factor (SCF) by stromal cells, either soluble or in a membrane-bound form, promotes survival and self-renewal through interaction with c-kit receptor^62^. Thus, we characterised the expression levels of key cytokines, signalling molecules, and adhesion proteins, by MSCs conditioning @BMN. Transcriptomic analyses indicated the stable expression of SCF, IL-6, Angiopoietin1 (Ang1), Thrombopoietin 1 (TPO-1), V-CAM, N-Cadherin (N-cad) and Jagged1 (Jag1) over a 15-day culture period (Figure 4A), suggesting that MSCs growing on bioemulsions and conditioning the @BMN maintain key cues able to regulate the long-term culture of HSPCs. Moreover, increased expression levels were observed at day 15 (compared to day 4) for Vascular Cell Adhesion Molecule-1 (VCAM-1) and C-X-C motif Chemokine 12 (CXCL12, also known as SDF-1α), two important regulators of HSPC fate (Figure 4B). Importantly, when comparing CXCL12 expression on bioemulsions to 2D TCP, a significant (nearly 20-fold) enhancement was observed (Figure 4B). The expression of CXCL12 was further confirmed by immunostaining and confocal microscopy, revealing apparent clusters of this cytokine, cytoplasmic as well as pericellular, suggesting its effective release (Figure 4C). Similarly, abundant production and release of SCF by stromal cells in @BMN was observed (Figure 4C), confirming the expression of this key cytokine.

**Figure 4.**
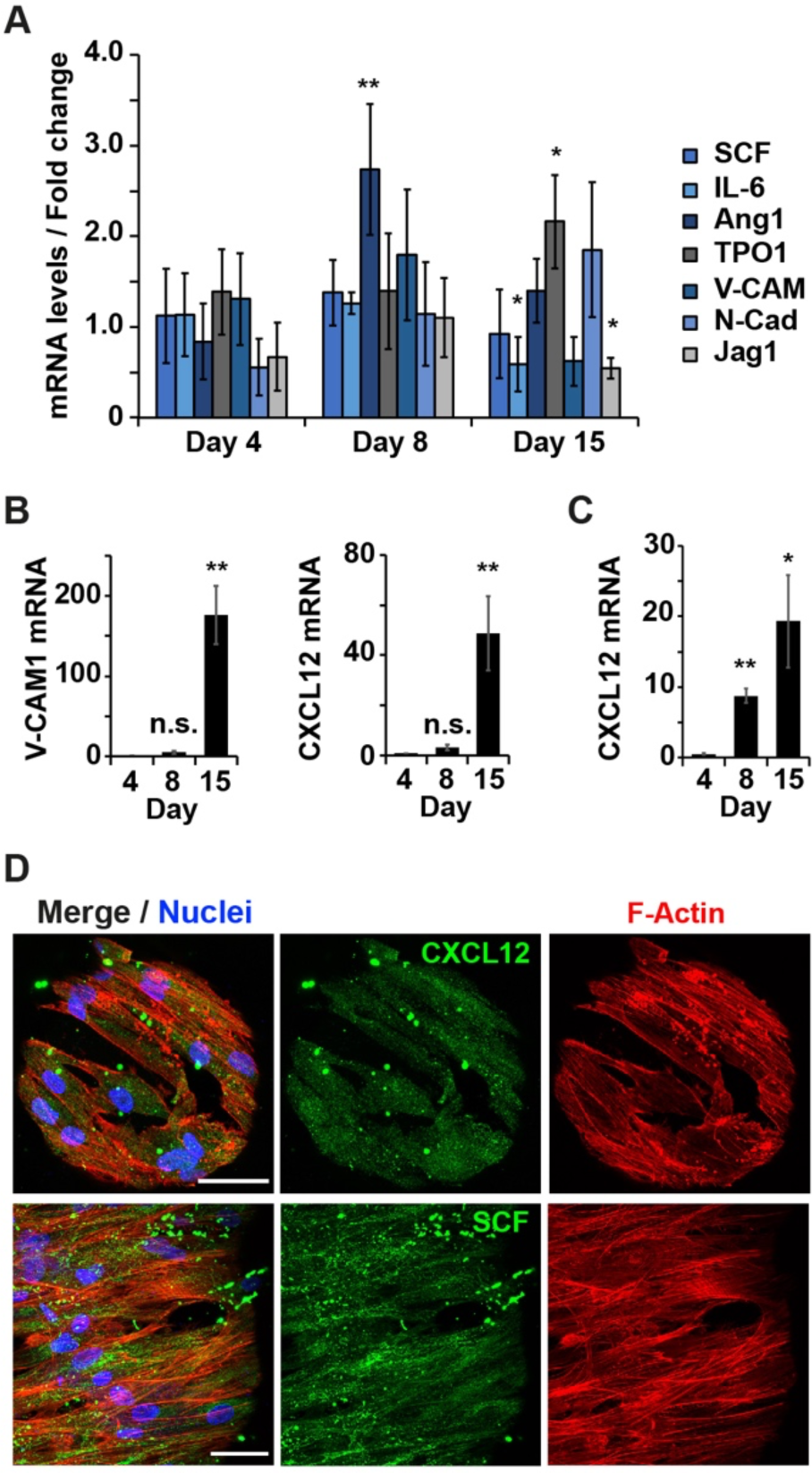
@BMN display secretory phenotypes priming these niches for HSPC implantation. A) Transcriptional quantification of cytokines and signalling molecules expressed by MSCs on bioemulsions, relative to levels measured on 2D TCP. Mean ± SD; N = 3; *, P<0.05, **P<0.01; statistical comparison is with the same condition and time point, but for cells cultured on 2D TCP. B) Quantification of mRNA levels of VCAM-1 (left) and CXCL12 (right) in MSCs cultured on bioemulsions relative to levels at day 4. Mean ±SD; N = 3; n.s., P>0.05; **, P<0.01. C) Quantification of mRNA levels of CXCL12 (right) in MSCs cultured on bioemulsions, relative to levels on 2D substrates at different timepoints. Mean ± SD; N = 3; n.s., P>0.05; **, P<0.01. D) z-Projection of confocal z-stacks of MSCs growing on biolemulsions, immunostained to visualize CXCL12(upper) and SCF (lower) production and secretion. Scale bar, 20 μm.

Overall, these results demonstrate that MSCs assembled a network of ECM proteins around and between microdroplets of bioemulsions, further supporting cell adhesion, and display a secretory phenotype associated with the regulation of HSPC fate decision. Together, this collection of biochemical and biophysical signals contributes to shaping a mature @BMN that may effectively maintain HSPC populations and may promote their expansion.

### Artificial BM microenvironments enable the expansion and maintenance of HSPCs

Having established @BMN recapitulating some of the key structural, mechanical, and biochemical components of the hematopoietic niche, we next examined their ability to support HSPC expansion. We first analysed how the maintenance of the CD34^+^/CD38^−^ populations is influenced by the presence of bioemulsions in mono-cultures. Interestingly, a slight increase in the percentage of CD34^+^/CD38^−^ cells was observed after 7 days of culture, but cell densities were comparable at day 14 (Supplementary Figure S6). Furthermore, HSPCs were growing in suspension in the presence of bioemulsions, with little interaction with the interface (Figure 5C). HSPCs were then seeded onto @BMN, after 7 days of conditioning by MSCs. Prior to HSPC seeding, MSC growth medium was replaced by HSPC medium (day 6, Figure 1C). To identify the specific impact of these MSC-primed niches on HSPC expansion, SCF alone (100 μM) was supplemented to SFEM II medium as a minimal formulation supporting HSPC culture^63^ (without the addition of any further cytokines) since it has been shown that SCF alone can enhance the density of cells with self-renewal capacity by 25%. This medium had no significant impact on the morphology and viability of MSCs themselves (Supplementary Figure S7). The interaction of HSPCs with MSCs, both on 2D TCP and in @BMN, was evidenced by time-lapse microscopy, using tagged HSPCs (Cell Tracker, Supplementary Videos S1 and S2). In addition, two-photon microscopy of co-cultures at day 15 (enabling deeper imaging) further revealed the presence of clusters of HSPCs and HSPC-derived cells within the core of the @BMN (Figures 5A and B, Supplementary video S3). Such clusters of HSPCs suggest that the combination of factors (geometry, mechanics and biochemical cues) within @BMN is conducive to high cycling and expansion of HSPCs on @BMN^64^. Indeed, this increased cycling was associated with a dramatic increase in HSPC proliferation when co-cultured with MSCs compared to HSPCs growing alone, in particular in @BMN (Figures 5 C and D). This correlates with the fact that the presence of SCF supplement alone was sufficient to induce the cycling and proliferation of HSPCs in co-culture but not on monoculture (Supplementary Figure S8). This observation highlights the strong impact that MSCs play in priming @BMN to promote HSPC expansion. Immunostaining and microscopy provided further evidence of the interactions between HSPCs and MSCs, with apparent internalisation of SCF and CXCL12 by HSPCs, suggesting a direct impact of the secretory phenotype of MSCs on HSPCs (Figure 5E). After 8 days in co-culture (day 15), HSPC proliferation rates remained high, while no MSC staining for Ki-67 was observed, further suggesting that MSCs remain quiescent at confluency, but support HSPC proliferation (Figures 5F and G).

**Figure 5.**
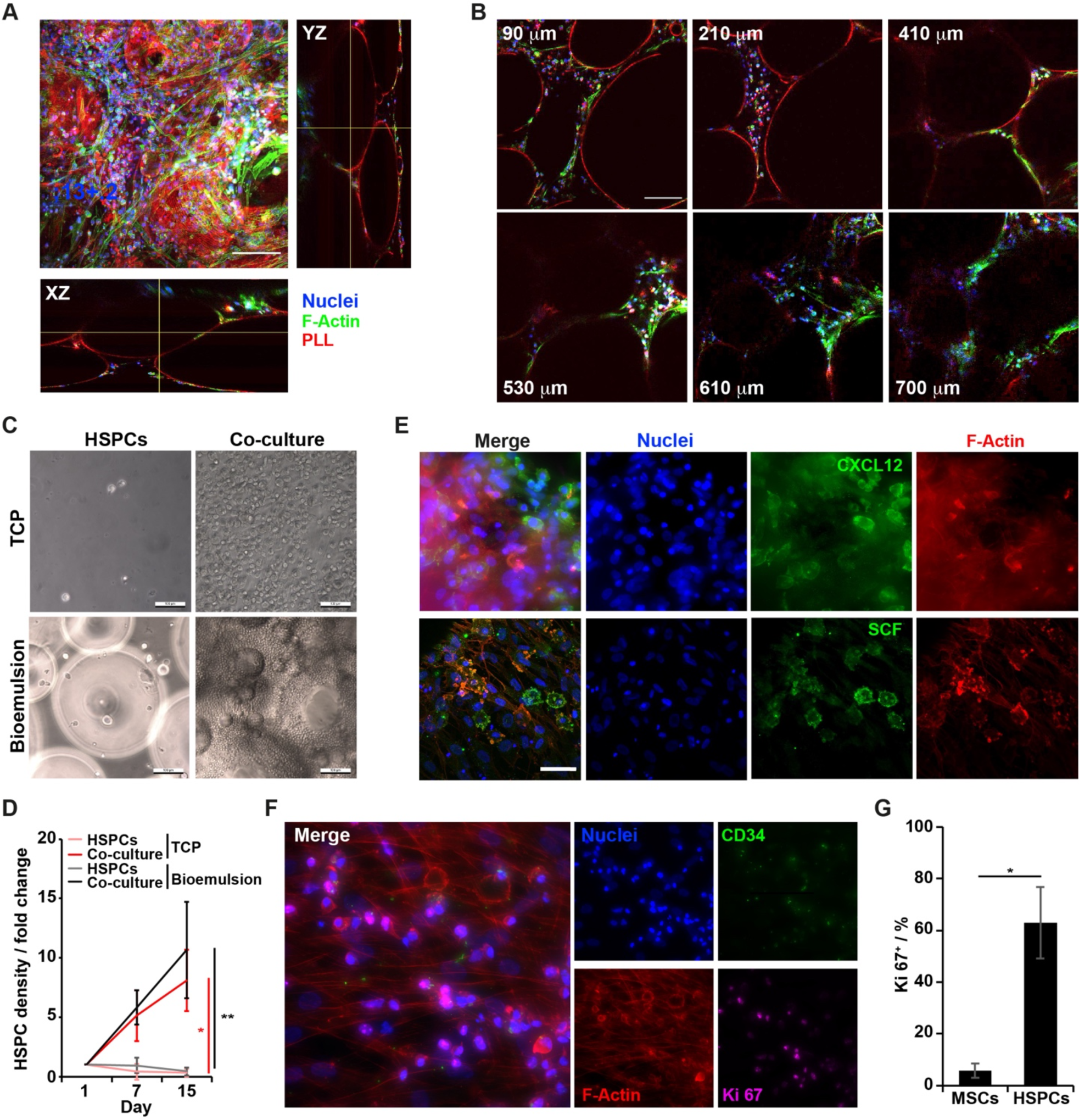
@BMN enable the expansion of HSPCs in vitro. A) z-Projection of two-photon confocal z-stacks of @BMN with HSPCs grown for 15 days, and respective XZ and YZ sections. Scale bar, 50μm. B) Selected confocal sections along the 700 μm z-stack. Scale bar, 50 μm. C) Brightfield images of HSPCs mono- and co-cultures with MSCs at day 7, on 2D TCP and on bioemulsions. Scale bar, 100 μm. D) Corresponding quantification of the evolution of HSPC densities at different time points. Mean ±SD; N = 3; *, P < 0.05, **, P < 0.01. E) z-Projections of confocal z-stacks of co-cultures growing on @BMN, immunostained to visualize the localisation of key cytokines secreted by MSCs (CXCL12 and SCF, green). Scale bar, 50 μm. F) z-Projection of confocal z-stack of co-cultures growing on @BMN, immunostained to evaluate cell cycling (Ki-67, green). Scale bar, 50 μm. G) Corresponding quantification of percentages of Ki-67^+^ cells. Mean ± SD; N = 3; *, P < 0.05.

The phenotype of HSPCs cultured in @BMN was next examined. HSPCs can be classified as short-term (ST)- and long-term (LT)-repopulating HSCs, based on their ability to repopulate the BM after transplantation^65^. While both cell types have the capacity to sustain haematopoiesis, the BM ST-HSCs can only do so for a short period of time. The LT-HSCs, however, have a long-term reconstituting capacity (>3-4 months) and are thus crucial to clinical success of associated therapies. Therefore, we investigated whether our @BMNs s supported the maintenance of an LT-HSCs population. LT-HSCs and ST-HSCs can be discriminated via their surface marker profiles. While both subpopulations are CD34^+^/CD38^−^/CD45RA^−^, LT-HSCs also express CD90. Flow cytometry analysis demonstrated that a small sub-population of CD34^+^/CD38^−^/CD45RA^−^/CD90+ cells was found in co-culture at days 7 and 15 but was absent when HSPCs where cultured alone (Figures 6A and B; Supplementary Figures S6C and S9). The presence of this LT-HSCs subpopulation, after 15 days of culture is particularly promising for the *in vitro* expansion of HSPCs in co-culture, with realistic therapeutic capacity. We next investigated the identity of the hematopoietic cells derived from the initial CD34^+^ that have proliferated in TCP co culture and @BMNs, compared to those derived from HSPCs growing in suspension. We identified hematopoietic cells including progenitor cells and more mature cells (CD45^+^), natural killer cells (NK, CD45^+^/CD56^+^), cells from the megakaryocytic lineage (CD41^+^), and red blood cells (CD45^−^/GlyA^+^). Given that most cells in the human hematopoietic system express the pan-leukocyte receptor CD45, we compared the proportion of cells expressing this marker in both conditions and found that this population was significantly more represented in co-culture (Supplementary Figure S9). This confirms the rapid loss of the hematopoietic identity of HSPCs grown in monoculture, in low cytokine media. The percentages of erythrocytes and megakaryocytes remain similar in both conditions indicating that their abundance is not affected by the culture system. However, there was a slight increase in differentiation towards the lymphoid lineage, as evidenced by a small but significant increase in densities of CD56^+^ cells (the archetypal marker for NK cells), in @BMN.

**Figure 6.**
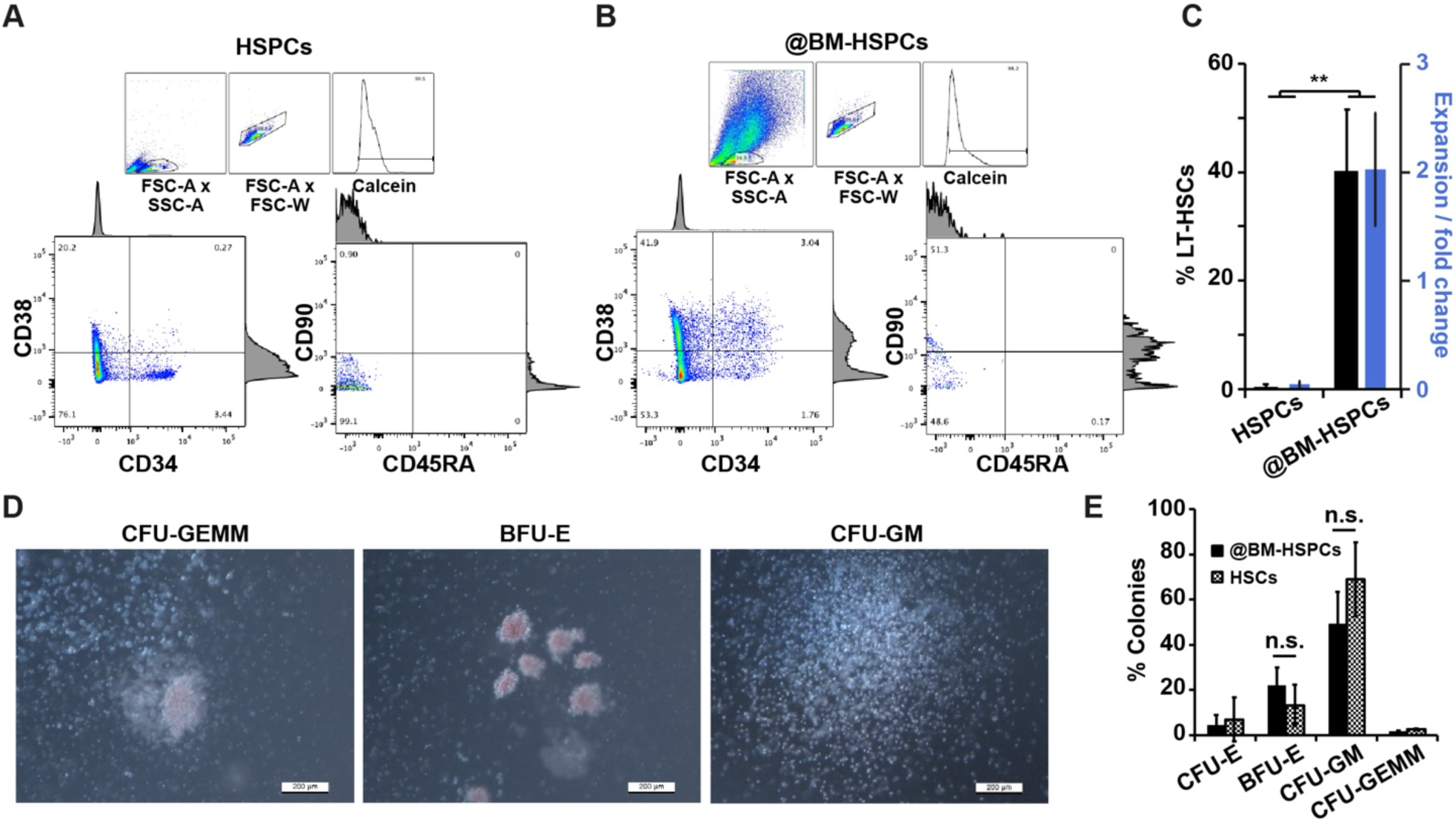
@BMN maintain the stemness of HSPCs *in vitro*. A and B) Gating strategy for the quantification of HPSC specific surface markers. After doublet exclusion, mononuclear cells were gated based on low/high expression of calcein to identify live cells and exclude bioemulsions and dead cells. CD34^+^/CD38^−^ subset was selected and CD90^+^/CD45RA^−^ cells were subsequently gated to identify the population of interest. C) Fold expansion of CD34^+^ cells and corresponding percentage of CD34^+^/CD38^−^/CD90^+^/CD45RA^−^ subpopulations of HSPCs, growing alone or within @BMN, recovered at day 15. Mean ± SD; N = 4; (**, P<0.01). D) Brightfield images of CFU-GEMM, BFU-E and CFU-GM colonies; Scale bar, 200 μm. E) Quantification of the densities of each type of colonies arising from the CFU-Assay, taken from CD34^+^ cells isolated from monocultures and @BMN. Mean ± SD; N = 3; *, P<0.05. No statistically significant differences were observed.

It has long been established that CD34^+^ cell are responsible of reconstituting haematopoiesis in transplanted patients, with higher CD34^+^ cell doses having a significant positive impact on clinical outcome^25,66^. *Ex vivo* selection of CD34^+^ for allogenic transplantation has also shown efficacy for the depletion of alloreactive donor T cells from grafts, translating into a reduction in the risk of developing acute and chronic graft-vs-host disease (GVHD)^67^. To assess the ability of @BMN to promote CD34^+^ population expansion, CD34^+^ cells were selected via magnetic assisted cell sorting and quantified. Whilst monocultures of HSPCs on bioemulsions were associated with a drastic reduction in the number of CD34^+^ cells, after 14 days of co-culture, the densities of CD34^+^ cells doubled (Figure 6C). This result is particularly striking, since no additional cytokines were supplemented, apart from basal SCF. In contrast, co-cultures typically require supplementation with additional relevant cytokines such as thrombopoietin and Flt-3^68^ in order to achieve such high populations of CD34^+^ HSPCs.

To further establish the retention of stemness and therapeutic potential of HSPCs grown on @BMN, we performed CFU assays on selected CD34^+^ cells isolated from long-term mono- and @BM co-cultures. Our results indicate that there were no significant changes in the multipotent capacity of CD34^+^ grown on the two conditions tested as the number of CFU-GEMM were similar (Figures 6D and E). Likewise, similar BFU-E, CFU-E and CFU-GM colony densities generated upon culture on @BMN as for CD34^+^ cells in monoculture were observed (Figure 6E). Moreover, the percentage of colonies obtained are within the normal ranges described for fresh CD34^+69^, indicating that long-term culture on @BMN has no negative impact in the clonogenic potential of CD34^+^ cells.

### Scaling up the expansion of HPSCs in artificial BM niche bioreactors

The implementation of *in vitro* co-culture systems for the expansion of HSPCs has so far struggled with appropriate substrate to support the growth of adherent MSCs whilst maintaining HSPC phenotype in a scalable format. Having demonstrated that bioemulsions provide a biomimetic BM microenvironment that supports the growth and maintenance of phenotype of HSPCs (Figures 5 and 6), the scalability of this platform was evaluated. To do so, @BMN formation, priming, and HPSC culture was scaled-up 100-fold, in a conical flask bioreactor format (Figures 7A and B) under agitation in an orbital shaker, comparable to systems used for the culture of planktonic cells, yeast or bacteria. Under these culture conditions, @BMN were still able to form (Figure 7B), providing biomimetic confinement and local mechanical properties. Viability assays performed on MSC long-term cultures (22 days) indicated no reduction in cell survival, in all conditions tested (2D TCP, static @BMN in 24 well-plate format and @BMN in bioreactor format; Figures 7C and D). Phenotypic analysis demonstrated the maintenance of MSC identity upon culture on bioemulsions within conical flask bioreactors, with high triple marker expression (CD73, CD90 and CD105; Supplementary Figure S10 A, B). Expression of fibronectin, CXCL12 and Nestin after 7 days of culture, right before seeding of HSPCs, was also confirmed by IF imaging (Supplementary Figure S11). After 15 days of co-culture, isolation of CD34^+^ cells was performed using magnetic cell sorting and cell numbers quantified. HSPC expansion in co-culture on conical flasks was found to be enhanced to 2.97 ± 0.3-fold, compared to HSPCs growing in monoculture 0.09 ± 0.13 times (Figure 7E). Thus, the expansion of HSPCs on @BMN in conical flasks was found to produce 33.4 times more CD34^+^ cells than the traditional TCP monoculture suspension method using a low-cytokine medium. To further characterise the CD34^+^ population expanded in conical flask bioreactors, the percentage of LT-HSCs was investigated to confirm the maintenance of a more abundant population compared to HSPC monocultures (Supplementary Figure S12). the multipotency of CD34^+^ HSPCs recovered was assessed via CFU assay (Figure 7F). This confirmed the maintenance in their clonogenic potential, with comparable levels of commitment to multilineage differentiation after long-term culture to that of HSPCs in TCP monoculture and fresh cells, suggesting an equal distribution of different progenitor types for the conditions tested. Therefore, the long-term co-culture of CD34^+^ cells on @BM niches did not affect their clonogenic potential and cells retained their myeloid differentiation potential. Altogether, these results demonstrate the scalability of bioemulsion-based @BMN, for the expansion of HSPCs *in vitro*, achieving a 33-fold increase in CD34^+^ cells compared to suspension monocultures, with addition of SCF alone, and a net ∼3-fold increase in HSPCs. This implies that realistic volumes of bioemulsions (e.g. 40 ml emulsion phase) could support the growth of considerable densities of HSPC-derived cells, yielding an expected 6 10^6^ CD34^+^ cells, an output well within the range of 2 to 8 10^6^ cells/kg CD34^+^, considered to be required for successful transplantation. Therefore @BMNs are attractive for the scalable *ex vivo* expansion of HSPCs prior to infusion.

**Figure 7.**
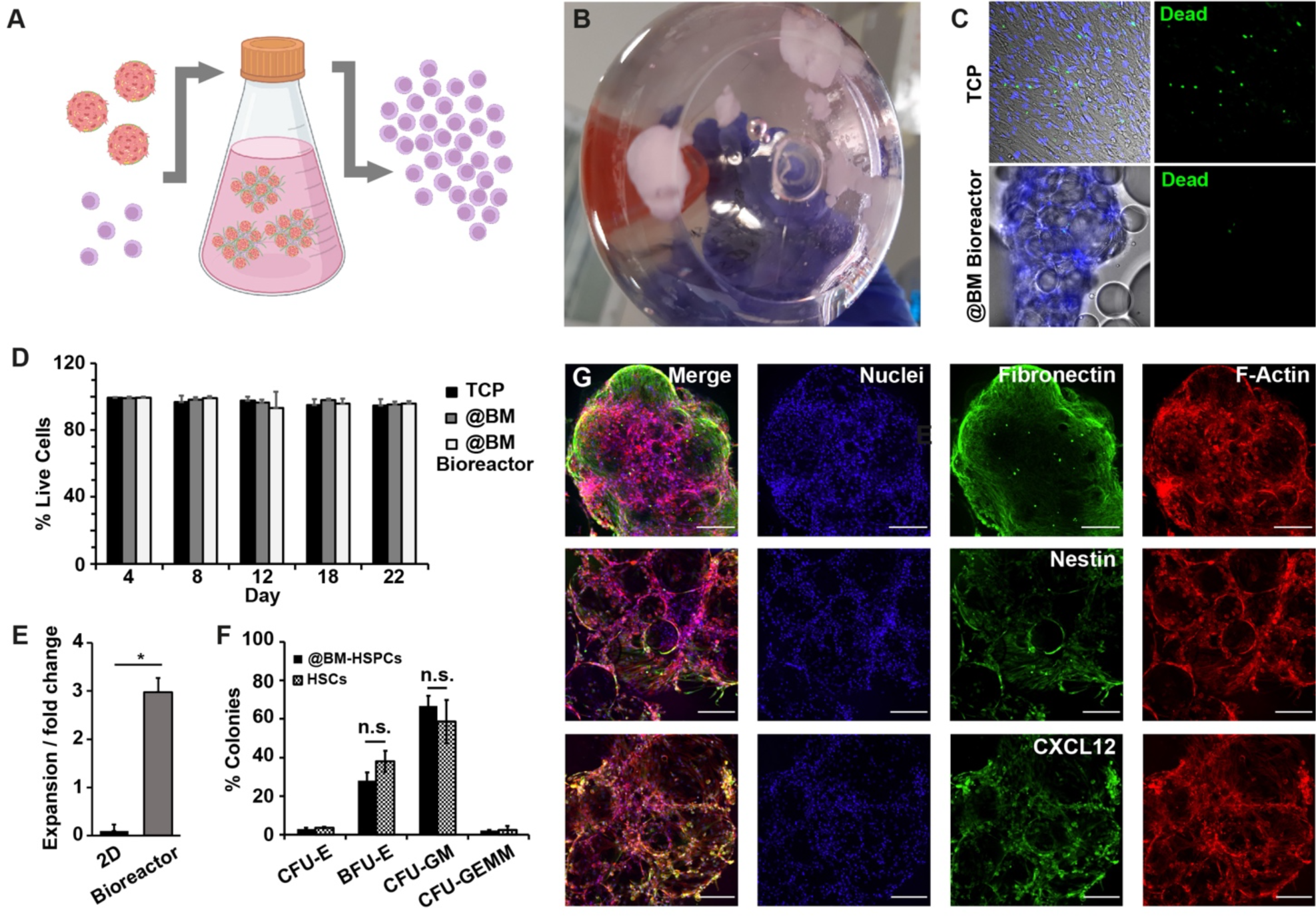
@BMN enable the scale up of *in vitro* expansion of HSPCs. A) Schematic representation of the scale-up of @BMN in a conical flask bioreactor format. B) Image of artificial BM microtissue formed during the culture of MSCs on bioemulsions, still in the culture conical flask. C) Hoechst/ethidium staining of MSCs growing on 2D and on @BMN, in multi-well plate and in conical flask bioreactors, after 22 days of culture in MSC medium. Scale bar, 100 µm. D) Corresponding quantification of percentages of live cells at days 4, 8 12, 18 and 22. Mean ± SD; N = 3; No statistically significant differences were observed between these conditions. E) Quantification of the expansion levels of HSPCs grown on @BMN in conical flask bioreactor, compared to monoculture in suspension. Mean ± SD; N = 3; *, P < 0.05. F) Quantification of the percentages of clonogenic differentiation observed in CFU assays carried out with CD34^+^ cells isolated from monoculture and co-culture in @BMN. Mean ± SD; N = 3; *, P < 0.05. G) z-Projection of confocal z-stack of co-cultured @BMN grown in conical flask bioreactor for 14 days. Scale bar, 100 μm.

## 3. Discussion

Conditions such as myeloid disorders, acute lymphoid disorders or autoimmune diseases often require high doses of chemotherapy or radiation exposure that severely compromise the homeostasis of the BM. These harsh procedures are typically followed by a HSPC transplant that aims to reconstitute a healthy hematopoietic system^1^. Current strategies for patients suffering from these disorders primarily consist of allogenic transplantations, utilizing stem cells from different sources such as umbilical cord in infants^70^, mPB^4^ or, less frequently, BM^71^. However, these procedures are often associated with slow hematopoietic recovery and a high risk of GvHD in allogenic transplantation^72^. Complementary strategies based on the *ex vivo* expansion of hematopoietic progenitors have been developed in the last few decades, and some of these approaches have already moved from pre-clinical into clinical trials^17,73–78^. However, the low cellular output associated with these procedures, need for high levels of cytokines, and the limited capacity of infused cells to enable neutrophil engraftment and platelet recovery, remain a critical hurdle to address.

Several studies have indicated the importance of the role of the BM niche in the regulation of HSPC phenotype and proliferation^7^. Therefore, the engineering of biomimetic @BMN is attractive to reproduce some of the structural, mechanical, and biochemical features of the native hematopoietic niche. A strategy often explored has proposed the development of BM models using engineered hydrogels or scaffolds, to confer a 3D structure with mechanical properties mimicking those of physiological bone marrow^16,79,80^. However, these systems typically lack the microstructure and mechanical anisotropy of the BM microenvironment and are difficult to scale-up (e.g., limited nutrient diffusion, poor processability and cell recovery). In addition, in most approaches, cell expansion is typically sustained by the addition of cytokines, with no or limited contact with other supporting stromal cells due to the constraints regarding scalability in co-culture. Indeed, the expansion of adherent cells, in particular MSCs of various origins, has benefited from the implementation of plastic microcarriers for culture in 3D bioreactors^81^, although they introduce further restrictions, regarding the processing of cell products^82,83^. To the best of our knowledge, these platforms have not been successfully applied to the co-culture and production of HSPCs with clinically relevant phenotypic potential. Instead, HSPCs have been co-cultured with stromal cells in immobilized in porous glass or ceramic structures in fixed-bed bioreactors, achieving 7- to 46-fold expansion levels, although relying on the administration of additional cytokine cocktails^16,84^. However, these systems are either requiring supplementation of complex growth factor cocktails or associated with bioprocessing protocols that are difficult to scale up.

In this context, bioemulsions appear promising strategies to recapitulate several key features of the BM niche while providing a scalable platform with continuous perfusion for cell expansion. Specifically, our data demonstrated the ability to capture the microstructure and local as well as mesoscale mechanical properties of the BM microenvironment (Figures 1 and 3), as well as some of the biochemical cues that regulate cycling and fate decision in HSPCs. Indeed, whereas the endosteum niche, enriched in fibronectin has been reported to display relatively high Young’s Moduli (near 40-50 kPa), the perivascular niche and central medullary regions are much softer with moduli in the range of 3 kPa and 0.3 kPa, respectively^28^. Although liquid substrates are fluids with mechanical properties incompatible with the establishment of cell adhesions and the regulation of integrin-mediated signalling pathways, the high interfacial shear moduli and elasticity associated with protein nanosheets stabilising microdroplets^26,27,37^ are proposed to capture the local mechanical properties of the cortical/membrane/pericellular matrix associated with adipocyte-rich tissues^33^. This results in local mechanical properties (from AFM indentation data, Figure 3) that correlate well between @BMN and native BM niches. In addition, the mechanical properties of protein nanosheets stabilising bioemulsion-based @BMN are sufficiently tough and elastic^27,37,40^ that they can resist cell-mediated contractile forces and enable the remodelling of a dense nanofibrous extra-cellular matrix recapitulating some of the compositional and topographical features of the BM matrix.

Although such observations are in good agreement with the recognised importance of matrix mechanics and viscoelasticity on cell adhesion and matrix remodelling^85–88^, the impact that such local mechanics may have on secretory phenotypes is less clear. Indeed, previous reports have drawn correlations between microenvironment mechanics, degradation, matrix deposition and microstructure and secretory phenotypes^89,90^, but the detailed molecular mechanisms regulating such behaviours remain unclear. Similarly, the observation that MSCs growing on bioemulsions overexpress two key cytokines associated with the regulation of HSPC phenotype, VCAM-1 and CXCL12 (Figure 3), cannot be at present solely attributed to one key physical or biochemical parameter associated with protein nanosheet interfaces and bioemulsions. But, however, it could be proposed that increased hypoxia and growth arrest at the core of the @BMN may be inducing a higher expression of CXCL12 in this platform compared to 2D configurations^91–93^. In turn, the combined impact of microstructure, matrix remodelling, and cytokine composition is proposed to capture the biochemical context associated with the BM microenvironment while promote the formation of gradients of cytokines and signal retention.

From a translational point of view, although the supplementation of defined cytokine and growth factor cocktails is attractive for the development of cell therapy products, it is increasingly recognised that the supplementation of cytokines to HSPC cultures not only induces cell proliferation but also boosts differentiation and, thus, a dramatic loss of CD34^+^ cells posing important hurdles to the translation of stem cell therapies. More recent approaches aiming to preserve the self-renewal potential of HSPCs *in vivo* are currently being tested. Among them, co-culture with supporting cells such as MSCs circumvents these pitfalls by, to some extent, preventing HSPC differentiation^17^. However, appropriate cell substrates enabling MSCs adhesion at scale need to be developed in order to reach relevant HSPC expansion levels in co-culture. While producing sufficient cell numbers using 2D platforms is extremely unrealistic as such approaches cannot be used routinely, the use of our bioemulsion technology in bioreactors could be a potential solution to this scalability challenge. Our data suggests that expansion of HSPCs in co-culture can be performed at scale by using bioemulsions while maintaining reconstituting potential of the CD34^+^ population. Furthermore, the bioemulsion system is easy-to-handle and circumvents the use of enzymatic treatment for cell harvesting^36^, which is not only costly but might cause unintended damage to cells and/or phenotypical changes. While we achieved an HSPC expansion of ∼3-fold using a low cytokine medium, our technology is compatible with the use of additional strategies to promote further expansion such as the use of additional cytokines^17,94^

In addition, a number of other compatible approaches have already demonstrated the capacity to enhance expansion *in vitro*, prior to re-implantation and are already being clinically tested. These include the use of novel small molecules such as StemRegenin-1, UM171 and nicotinamide or the induction of Notch signaling pathways among others. Other innovative strategies consist of the use of genetic engineered bacteria expressing relevant cytokines and adhesion molecules to prime a hematopoietic niche *in vitro*^95^. Finally, the influence of changes in oxygen content within the niche, on HSPC self-renewal potential should not be minimised^96,97^, and strategies enabling the regulation of oxygen tension in bioreactors could help further enhance the scale-up of HSPCs production^98^.

Overall, biomimetic @BMN offer unique opportunities for the scale-up of HPSC expansion in 3D bioreactors. This work not only demonstrates the feasibility of this concept, but also calls for further studies completing our understanding of the impact of biophysical and biochemical factors, and their crosstalk on HSPC fate decision. The associated niches and engineered microenvironments may not only find application for the transplantation of stem cells for regenerative medicine, but also for the design of advanced *in vitro* models for the testing of safety and efficacy of therapeutics. In addition, this report further establishes the potential of bioemulsions to replace conventional solid microcarriers, based of plastics or hydrogels, presenting a hurdle to cell manufacturing in terms of processing, microplastic contamination and production costs. Indeed, new bioemulsion systems based on plant-based oils or biomedical grade oils^99^ offer alternative to these issues and offer opportunities for the direct delivery of cell colonies, without requirement for further cell separation or processing.

## 4. Experimental Section

### Preparation of bioemulsions

Bioemulsions were generated by mixing the fluorinated oil (Novec 7500, ACOTA) supplemented with 10 µg/ml of Pentafluorobenzoyl chloride (Sigma-Aldrich) with the protein aqueous solution (1 mg/ml of PLL, Sigma-Aldrich, in PBS pH 10.4), at a 1:2 ratio. The vial was shaken for 15 s and incubated for 1 h at room temperature. The aqueous phase was aspirated, and the emulsions thoroughly washed with PBS to remove excess PLL. Emulsions were then incubated with Fibronectin (20 μg/ml, Millipore, pH 7.4) for 1 h at room temperature. For cell culture, 50 µl of emulsions/well were transferred 24-well plates treated with 25 μg/ml poly(L-lysine)-graft-polyethylene glycol (PLL-g-PEG, SuSoS).

### Evaluation of emulsion stability

The stability of the bioemulsions was assessed at different timepoints using Mastersizer 2000 according to manufacturer’s instructions. Particle size distribution was determined for each condition at the day of bioemulsion formation (Day 1) and after 7 days and 15 days kept at 37 °C under agitation in an orbital shaker (60rpm). The emulsion stability was also monitored via bright-field microscopy.

### Mesenchymal stem cells (MSCs) culture and seeding on emulsions

Bone marrow-derived human mesenchymal stem cells were purchased from PromoCell and cultured in MSC Growth Medium 2 (PromoCell). MSCs between passages 2 and 6 were used for the experiments. For cell seeding on emulsion, MSCs were harvested using Accutase (PromoCell) and centrifuged at (200xg) for 5 min. 3×104 cells were seeded in 24-well plates containing 50 µl of PLL emulsion and 750 μl of MSC Growth Medium and cultured for 7 days in a humidified atmosphere with 5%CO2. For the 2D controls, 7,000 cells/well were seeded in 24-well plates containing 500 μl of MSC Growth Medium. Half of the medium was replaced with fresh medium every two days.

### Co-culture

Bone marrow-derived human CD34+ stem cells were purchased from Stemcell technologies and cultured in SFEM II medium (Stemcell technologies) containing 100 μM SCF (Stemcell technologies) according to manufacturer’s instructions. For co-culture, the medium on the 24-well plates containing the 7-day MSCs cultures was replaced with the SFEM II medium 1 day prior to CD34+ cell seeding. For co-culture, both on 2D and on emulsion, 5,000 cells were seeded in each well containing the MSC culture. For the HSPC control, cells were seeded on 2D or emulsions without previous addition of MSCs. Cultures of MSCs where no HSPCS were added were kept as the MSC-only controls. Cultures were kept for long-term experiments for a further 14 days at 37 °C in a humidified atmosphere with 5%CO2. Half of the medium was replaced with fresh medium every two days.

### Conical flask bioreactor scale-up

Scale up of the co-culture system was performed on 250 ml polycarbonate Erlenmeyer flasks (Corning) with Vent caps, using a total amount of medium of 100 ml. Volumes and amounts of further reagents and cells used where increased 100 times with respect to those in the culture on 24 well-plates. 10 ml of emulsions were used per conical flask while 3×106 MSCs and 5×105 CD34+ cells were seeded at the appropriate timepoints to mirror those on the 24-well plate. Cultures were kept under gentle agitation of 60 rpm on an orbital shaker, placed inside an incubator at 37 °C in a humidified atmosphere with 5% CO2

### Immunofluorescence staining

Cells in 2D (over borosilicate coverslips) and emulsions were fixed with 4% PFA for 20min and then permeabilised with 0.2% Triton X-100 for 10 min at 4 °C. Samples were then blocked with 3%BSA at room temperature for 30min. Primary antibodies were diluted in 3% BSA and incubated from 1h at 37 °C to overnight at 4 °C at the recommended concentrations depending on the antibody. After washing, secondary antibodies (Alexa Fluor488/647-conjugated, Thermofisher, 1:1000), DAPI (Life Technologies, 1:1000) and phalloidin-TRITC (Sigma-Aldrich, 1:500) were incubated for 1 hour in the dark and then washed with PBS. Cells cultured as 2D monolayers were mounted with fluoromount (Thermo Fisher Scientific) over glass slides. Cells cultivated on emulsions were transferred to μ-slide 8 wells plates (Ibidi) and preserved with PBS+0.05% Azide (Sigma-Aldrich) at 4 °C for a maximum period of 2 weeks. Primary antibodies used were: anti-Vinculin (#V9264, Sigma-Aldrich, 1:500) anti-Nestin (#ab22035, Abcam, 1:250) anti-K-i67 (#RM-9106, Epredia, 1:500), anti-Fibronectin (#F3648, Sigma-Aldrich, 1:500), anti-Laminin (#L9393, Millipore, 1:500), anti-Collagen I (#ab90395, Abcam, 1:1000), anti-CXCL12 (#ab9797, Abcam, 1:100), anti SCF(#ab64677, Abcam, 1:200), anti CD34-AlexaFluor488 (#60013AD, Stemcell technologies, 1:100).

### Microscopy

Zeiss LSM710 laser scanning microscope and Nikon CSU-W1 Spinning disk microscope with Two Photometrics Prime BSI sCMOS were used for confocal imaging. Objectives used were usually EC Plan-Neofluar20x/0.5 M27 and Plan-Apochromat 63×/1.4 Oil DIC M27 on the Zeiss LSM710. Objectives used in the Nikon CSU-W1 were CFI Plan Apochromat VC 20X Air and CFI Apochromat TIRF 60XC Oil. Widefield fluorescent images were taken using Leica DMi8 fluorescence microscope coupled to a Leica DFC9000 GT sCMOS camera. Objectives used were HC PL FLUOTAR L 20x/0.40 CORR PH1 and HC PL FLUOTAR L 40x/0.60 CORR PH2. For live-cell imaging, HSPCs where stained with Cell tracker CM-dil prior to seeding on the substrate (TCP or bioemulsions) containing the MSC feeder layer. Live imaging experiments were performed using incubator chamber accessories for each system at 37 °C and 5% CO2. Imaging medium was phenol red-free complete DMEM supplemented with 10% FBS and 25 mM Hepes. Images were acquired with a Lumascope 720 using Olympus LCAch N 20x/0.40 objective. Image and video analyses, z-stack projections and 3D deconvolution were done with ImageJ software.

### Viability staining and cell counting

Cell viability in the conical flask bioreactor format was assessed using Live/dead Viability/Cytotoxicity Kit (Invitrogen). A 300 ul sample of MSC emulsions was taken from the Erlenmeyer flask at different timepoints and placed in a 24-well containing a pre-warmed solution of PBS with Hoechst (1 mg/ml Thermofisher Scientific) and the LIVE/DEAD staining solutions. After 30 min incubation, cells were imaged using a Leica DMi8 epifluorescence microscope. Viability was compared to that of cells growing on monolayers at the same timepoints.

### Cryo-SEM

Cryo-SEM imaging was performed using a FEI Quanta 3D Dual Beam SEM. Bioemulsions, bovine BM tissues and @BM microtissue samples were collected at appropriate time points and mounted with O.C.T (Tissue-Tek) onto an aluminium stub on the specimen holder. Sample freezing was carride out via N2 plunging and frozen samples were transferred under vacuum to the cryo-preparation chamber attached to the microscope. Cryo-preparation was performed in a Gatan Alto 2500 cryogenic preparation cold stage, where samples was cut with a cold knife to uncover inner parts of the structure. The temperature of the sample was then raised to −125°C and passed through the transfer lock to the FEI Quanta 3D Dual Beam SEM cryo-stage, which was held at −125°C. The sample was sputter-coated using a platinum gas injection system. Imaging was performed using an accelerating voltage of 3 kV to 10 kV.

### Rheology

Rheological measurements were carried out using a hybrid rheometer (DHR-3) from TA Instruments. Samples were placed between an upper parallel standard Peltier plate geometry (20 mm) and the rheometer plate pre-heated at 37 °C. A gap of 500 μm was used for emulsions and 2 mm for the bovine bone marrow. After the geometry was lowered to the desired gap, the excess emulsion or bone marrow was trimmed with a spatula. First, frequency sweeps were carried out, recording the storage modulus (G’) and loss modulus (G’’) as a function of frequency in a range of 0.1-100 Hz, at 1% strain. Amplitude sweeps from 0.1 to 100 % strain were subsequently carried out, at an oscillating frequency of 1 rad/s.

### Atomic Force Microscopy

To perform AFM nanoindentation, samples were placed in a 35-mm petri dish and submerged in PBS. Force–distance measurements were carried out on a JPK NanoWizard 4 (JPK Instruments), with samples submerged in PBS at room temperature. Pyrex-nitride colloidal probes (diameter 15 µm; CP-PNPL-SiO-E-5, NanoAndMore) were used (spring constant (K) was calibrated, 0.08 N/m). Scans of 10 × 10 μm grid of indentation at three different locations across the surface of two different samples were acquired for each condition, and force-displacement curves were recorded. Indentations were carried out with a relative setpoint force of 2 nN and a loading rate of 5 μm s^−1^. Data were collected using JPK proprietary SPM software (JPK Instruments) and Young’s moduli were calculated using the Hertz model for a spherical tip.

### Flow cytometry

Single cell suspensions of cells growing on emulsions and on TCP were obtained via Accutase treatment and agitation. Cells were then washed in 0.1% BSA and centrifuged (300xg) for 10 min at 4 °C. Cells were stained for 45 minutes at RT using fluorescence-labelled antibodies and Calcein Violet (#C34858, Thermo Fisher Scientific) for cell viability. Labelled cells were then washed in PBS + 1% BSA and analyzed on a FACSCanto II flow cytometer (BD Biosciences). Data analysis was performed using FlowJo software (Tree Star). Conjugated antibodies used were: FITC anti-CD73 (#MCA6068F, Bio-Rad), APC anti-CD90 (#MCA90APC, Bio-Rad), PerCP/Cy5 anti-CD105 (#323216, BioLegend), FITC anti-CD38 (#356610, BioLegend), PE anti-CD34 (#343506, BioLegend), PE/Cy5 anti-CD45 (#304010,) APC anti-CD45RA (#304112, BioLegend), APC anti-CD41 (#303709, BioLegend), BV421 anti-CD56 (#362552, BioLegend) and PE anti-CD235a (#15805948, Fisher Scientific).

### qPCR

For Real-time quantitative PCR (qPCR) assay MSCs were grown either on 2D in 24-multiwell plates (7000 cells/well), or on emulsions (50μl emulsion/well 3×104/well). Total RNA was isolated at 4, 7 and 15 days and purified using RNeasy kit (Qiagen). 1 μg of RNA per condition was used to retrotranscribe into cDNA QuantiTect® Reverse Transcription Kit (Qiagen). qPCR was performed in 386 wells plates in QuantStudio™ Real-Time PCR System (Applied Biosystems™) using the Taqman® Gene Expression Assay kit following manufacturer’s instructions. Every condition was tested by 3 experimental replicates per gene. Experiments were conducted for an n = 3–4. BM2 (Hs00187842_m1, Thermo Fisher) gene was used for normalization. Relative quantification analysis was used to determine the fold change in RNA levels on emulsions either relative to early timepoints or 2D conditions using the Pfaffl model^100^. Details of Taqman probes used for target genes are listed in the supplementary information.

### CFU-Assay

The 1.0 × 10^3^ cells were harvested from each status after 7 days of expansion. Then, these cells were used in 3 mL of methylcellulose medium (H4435; Stem Cell Technology, Canada) plated in a 35-mm culture dish (Grainger). The cell’s dish with two uncovered dishes containing 3 to 4 mL sterile water was put into a 100-mm culture dish and covered. The sterile water dish was served to steadfastly keep up the humidity essential for colony development. H4435 contains 1% methylcellulose in Iscove’s modified Dulbecco’s medium (IMDM) (Sigma-Aldrich, Germany). After 14 days incubation at 37°C and 5% CO2, colony-forming units-granulocyte-macrophage progenitor (CFU-GM), granulocyte CFU (CFU-G), macrophage CFU (CFU-M), and granulocyte/erythrocyte/macrophage/megakaryocyte (CFU-GEMM); the multilineage HSPCs) colonies were counted. Colonies consisting of greater than, or equal to 50 cells were scored under an inverted microscope (Olympus IX81; Olympus).

## Supporting Information

Supporting Information is available from the author.

## Acknowledgements

Funding for this work from the European Research Council (ProLiCell, 772462 and ProBioFac, 966740) is gratefully acknowledged.

## References

1. THOMAS ED, LOCHTE HL, LU WC, FERREBEE JW. Intravenous infusion of bone marrow in patients receiving radiation and chemotherapy. N Engl J Med. 1957. doi:10.1056/NEJM195709122571102

2. Chabannon C, Kubal J, Bondanza A, et al. Hematopoietic stem cell transplantation in its 60s: A platform for cellular therapies. Sci Transl Med. 2018;10(436). doi:10.1126/scitranslmed.aap9630

3. Horan JT, Logan BR, Agovi-Johnson MA, et al. Reducing the risk for transplantation-related mortality after allogeneic hematopoietic cell transplantation: How much progress has been made? J Clin Oncol. 2011;29(7). doi:10.1200/JCO.2010.32.5001

4. Körbling M, Freireich EJ. Twenty-five years of peripheral blood stem cell transplantation. Blood. 2011;117(24). doi:10.1182/blood-2010-12-322214

5. Remberger M, Törlén J, Ringdén O, et al. Effect of total nucleated and CD34+ cell dose on outcome after allogeneic hematopoietic stem cell transplantation. Biol Blood Marrow Transplant. 2015;21(5). doi:10.1016/j.bbmt.2015.01.025

6. Dahlberg A, Delaney C, Bernstein ID. Ex vivo expansion of human hematopoietic stem and progenitor cells. Blood. 2011. doi:10.1182/blood-2011-01-283606

7. Morrison SJ, Scadden DT. The bone marrow niche for haematopoietic stem cells. Nature. 2014. doi:10.1038/nature12984

8. Ceafalan LC, Enciu AM, Fertig TE, et al. Heterocellular molecular contacts in the mammalian stem cell niche. Eur J Cell Biol. 2018. doi:10.1016/j.ejcb.2018.07.001

9. Tikhonova AN, Dolgalev I, Hu H, et al. The bone marrow microenvironment at single-cell resolution. Nature. 2019. doi:10.1038/s41586-019-1104-8

10. Sakurai M, Ishitsuka K, Ito R, et al. Chemically defined cytokine-free expansion of human haematopoietic stem cells. Nature. 2023;615(7950). doi:10.1038/s41586-023-05739-9

11. Shpall EJ, Quinones R, Giller R, et al. Transplantation of ex vivo expanded cord blood. Biol Blood Marrow Transplant. 2002. doi:10.1053/bbmt.2002.v8.pm12171483

12. Verovskaya E V., Dellorusso P V., Passegué E. Losing Sense of Self and Surroundings: Hematopoietic Stem Cell Aging and Leukemic Transformation. Trends Mol Med. 2019. doi:10.1016/j.molmed.2019.04.006

13. Bello AB, Park H, Lee SH. Current approaches in biomaterial-based hematopoietic stem cell niches. Acta Biomater. 2018. doi:10.1016/j.actbio.2018.03.028

14. Chramiec A, Vunjak-Novakovic G. Tissue engineered models of healthy and malignant human bone marrow. Adv Drug Deliv Rev. 2019. doi:10.1016/j.addr.2019.04.003

15. Xiao Y, McGuinness CAS, Doherty-Boyd WS, Salmeron-Sanchez M, Donnelly H, Dalby MJ. Current insights into the bone marrow niche: From biology in vivo to bioengineering ex vivo. Biomaterials. 2022;286. doi:10.1016/j.biomaterials.2022.121568

16. Bourgine PE, Klein T, Paczulla AM, et al. In vitro biomimetic engineering of a human hematopoietic niche with functional properties. Proc Natl Acad Sci U S A. 2018. doi:10.1073/pnas.1805440115

17. De Lima M, McNiece I, Robinson SN, et al. Cord-blood engraftment with ex vivo mesenchymal-cell coculture. N Engl J Med. 2012. doi:10.1056/NEJMoa1207285

18. Méndez-Ferrer S, Michurina T V., Ferraro F, et al. Mesenchymal and haematopoietic stem cells form a unique bone marrow niche. Nature. 2010. doi:10.1038/nature09262

19. Donnelly H, Ross E, Xiao Y, et al. Bioengineered niches that recreate physiological extracellular matrix organisation to support long-term haematopoietic stem cells. bioRxiv. 2022.

20. Ramalingam P, Butler J, Poulos M. Endothelial mTOR Preserves Hematopoietic Stem Cell Fitness By Suppressing Thrombospondin1. Blood. 2022;140(Supplement 1). doi:10.1182/blood-2022-162766

21. Luo Y, Shao L, Chang J, et al. M1 and M2 macrophages differentially regulate hematopoietic stem cell self-renewal and ex vivo expansion. Blood Adv. 2018. doi:10.1182/bloodadvances.2018015685

22. Ingavle G, Vaidya A, Kale V. Constructing 3D microenvironments using engineered biomaterials for HSC expansion. Tissue Eng Part B Rev. 2019. doi:10.1089/ten.TEB.2018.0286

23. Lee HJ, Li N, Evans SM, Diaz MF, Wenzel PL. Biomechanical force in blood development: Extrinsic physical cues drive pro-hematopoietic signaling. Differentiation. 2013. doi:10.1016/j.diff.2013.06.004

24. Schmal O, Seifert J, Schäffer TE, Walter CB, Aicher WK, Klein G. Hematopoietic stem and progenitor cell expansion in contact with mesenchymal stromal cells in a hanging drop model uncovers disadvantages of 3D culture. Stem Cells Int. 2016. doi:10.1155/2016/4148093

25. Ringdén O, John Barrett A, Zhang MJ, et al. Decreased treatment failure in recipients of HLA-identical bone marrow or peripheral blood stem cell transplants with high CD34 cell doses. Br J Haematol. 2003;121(6). doi:10.1046/j.1365-2141.2003.04364.x

26. Kong D, Megone W, Nguyen KDQ, Di Cio S, Ramstedt M, Gautrot JE. Protein Nanosheet Mechanics Controls Cell Adhesion and Expansion on Low-Viscosity Liquids. Nano Lett. 2018;18(3). doi:10.1021/acs.nanolett.7b05339

27. Kong D, Peng L, Bosch-Fortea M, et al. Impact of the multiscale viscoelasticity of quasi-2D self-assembled protein networks on stem cell expansion at liquid interfaces. Biomaterials. 2022;284:121494. doi:10.1016/j.biomaterials.2022.121494

28. Jansen LE, Birch NP, Schiffman JD, Crosby AJ, Peyton SR. Mechanics of intact bone marrow. J Mech Behav Biomed Mater. 2015;50. doi:10.1016/j.jmbbm.2015.06.023

29. Zhou BO, Yu H, Yue R, et al. Bone marrow adipocytes promote the regeneration of stem cells and haematopoiesis by secreting SCF. Nat Cell Biol. 2017;19(8). doi:10.1038/ncb3570

30. Naveiras O, Nardi V, Wenzel PL, Hauschka P V., Fahey F, Daley GQ. Bone-marrow adipocytes as negative regulators of the haematopoietic microenvironment. Nature. 2009;460(7252). doi:10.1038/nature08099

31. Fairfield H, Falank C, Farrell M, et al. Development of a 3D bone marrow adipose tissue model. Bone. 2019. doi:10.1016/j.bone.2018.01.023

32. Müller E, Grinenko T, Pompe T, Waskow C, Werner C. Space constraints govern fate of hematopoietic stem and progenitor cells invitro. Biomaterials. 2015. doi:10.1016/j.biomaterials.2015.02.095

33. Alkhouli N, Mansfield J, Green E, et al. The mechanical properties of human adipose tissues and their relationships to the structure and composition of the extracellular matrix. Am J Physiol Metab. 2013;305(12):E1427–E1435. doi:10.1152/ajpendo.00111.2013

34. Peled T, Shoham H, Aschengrau D, et al. Nicotinamide, a SIRT1 inhibitor, inhibits differentiation and facilitates expansion of hematopoietic progenitor cells with enhanced bone marrow homing and engraftment. Exp Hematol. 2012. doi:10.1016/j.exphem.2011.12.005

35. Crane GM, Jeffery E, Morrison SJ. Adult haematopoietic stem cell niches. Nat Rev Immunol. 2017. doi:10.1038/nri.2017.53

36. Peng L, Gautrot JE. Long term expansion profile of mesenchymal stromal cells at protein nanosheet-stabilised bioemulsions for next generation cell culture microcarriers. Mater Today Bio. 2021;12. doi:10.1016/j.mtbio.2021.100159

37. Kong D, Peng L, Di Cio S, Novak P, Gautrot JE. Stem Cell Expansion and Fate Decision on Liquid Substrates Are Regulated by Self-Assembled Nanosheets. ACS Nano. 2018. doi:10.1021/acsnano.8b03865

38. Megone W, Kong D, Peng L, Gautrot JE. Extreme reversal in mechanical anisotropy in liquid-liquid interfaces reinforced with self-assembled protein nanosheets. J Colloid Interface Sci. 2021;594. doi:10.1016/j.jcis.2021.03.055

39. Omatsu Y, Sugiyama T, Kohara H, et al. The Essential Functions of Adipo-osteogenic Progenitors as the Hematopoietic Stem and Progenitor Cell Niche. Immunity. 2010. doi:10.1016/j.immuni.2010.08.017

40. Peng L, Matellan C, Bosch-Fortea M, et al. Mesenchymal Stem Cells Sense the Toughness of Nanomaterials and Interfaces. Adv Healthc Mater. 2023. doi:10.1002/adhm.202203297

41. Isern J, Méndez-Ferrer S. Stem cell interactions in a bone marrow niche. Curr Osteoporos Rep. 2011. doi:10.1007/s11914-011-0075-y

42. Eagle H, Levine EM. Growth regulatory effects of cellular interaction. Nature. 1967. doi:10.1038/2131102a0

43. Cao H, Heazlewood SY, Williams B, et al. The role of CD44 in fetal and adult hematopoietic stem cell regulation. Haematologica. 2016. doi:10.3324/haematol.2015.135921

44. Choi JS, Harley BAC. Marrow-inspired matrix cues rapidly affect early fate decisions of hematopoietic stem and progenitor cells. Sci Adv. 2017. doi:10.1126/sciadv.1600455

45. Rodgers KD, San Antonio JD, Jacenko O. Heparan sulfate proteoglycans: A GAGgle of skeletal-hematopoietic regulators. Dev Dyn. 2008;237(10). doi:10.1002/dvdy.21593

46. Ahmed M, ffrench-Constant C. Extracellular Matrix Regulation of Stem Cell Behavior. Curr Stem Cell Reports. 2016. doi:10.1007/s40778-016-0056-2

47. Wasnik S, Kantipudi S, Kirkland MA, Pande G. Enhanced Ex Vivo Expansion of Human Hematopoietic Progenitors on Native and Spin Coated Acellular Matrices Prepared from Bone Marrow Stromal Cells. Stem Cells Int. 2016. doi:10.1155/2016/7231567

48. Dao MA, Hashino K, Kato I, Nolta JA. Adhesion to fibronectin maintains regenerative capacity during ex vivo culture and transduction of human hematopoietic stem and progenitor cells. Blood. 1998. doi:10.1182/blood.v92.12.4612.424k04_4612_4621

49. Yokota T, Oritani K, Mitsui H, et al. Growth-supporting activities of fibronectin on hematopoietic stem/progenitor cells in vitro and in vivo: Structural requirement for fibronectin activities of CS1 and cell-binding domains. Blood. 1998. doi:10.1182/blood.v91.9.3263.3263_3263_3272

50. Susek KH, Korpos E, Huppert J, et al. Bone marrow laminins influence hematopoietic stem and progenitor cell cycling and homing to the bone marrow. Matrix Biol. 2018. doi:10.1016/j.matbio.2018.01.007

51. Oswald J, Steudel C, Salchert K, et al. Gene-Expression Profiling of CD34 + Hematopoietic Cells Expanded in a Collagen I Matrix. Stem Cells. 2006. doi:10.1634/stemcells.2005-0276

52. Jiang J, Papoutsakis ET. Stem-Cell Niche Based Comparative Analysis of Chemical and Nano-mechanical Material Properties Impacting Ex Vivo Expansion and Differentiation of Hematopoietic and Mesenchymal Stem Cells. Adv Healthc Mater. 2013. doi:10.1002/adhm.201200169

53. Holst J, Watson S, Lord MS, et al. Substrate elasticity provides mechanical signals for the expansion of hemopoietic stem and progenitor cells. Nat Biotechnol. 2010. doi:10.1038/nbt.1687

54. Zhang X, Cao D, Xu L, et al. Harnessing matrix stiffness to engineer a bone marrow niche for hematopoietic stem cell rejuvenation. Cell Stem Cell. 2023;30(4):378–395.e8. doi:10.1016/j.stem.2023.03.005

55. Worzakowska M. Thermal and mechanical properties of polystyrene modified with esters derivatives of 3-phenylprop-2-en-1-ol. J Therm Anal Calorim. 2015;121(1). doi:10.1007/s10973-015-4547-7

56. Megone W, Roohpour N, Gautrot JE. Impact of surface adhesion and sample heterogeneity on the multiscale mechanical characterisation of soft biomaterials. Sci Rep. 2018;8(1). doi:10.1038/s41598-018-24671-x

57. Greenbaum A, Hsu YMS, Day RB, et al. CXCL12 in early mesenchymal progenitors is required for haematopoietic stem-cell maintenance. Nature. 2013. doi:10.1038/nature11926

58. Sugiyama T, Kohara H, Noda M, Nagasawa T. Maintenance of the Hematopoietic Stem Cell Pool by CXCL12-CXCR4 Chemokine Signaling in Bone Marrow Stromal Cell Niches. Immunity. 2006. doi:10.1016/j.immuni.2006.10.016

59. Simmons PJ, Masinovsky B, Longenecker BM, Berenson R, Torok-Storb B, Gallatin WM. Vascular cell adhesion molecule-1 expressed by bone marrow stromal cells mediates the binding of hematopoietic progenitor cells. Blood. 1992. doi:10.1182/blood.v80.2.388.bloodjournal802388

60. Frenette PS, Subbarao S, Mazo IB, Von Andrian UH, Wagner DD. Endothelial selectins and vascular cell adhesion molecule-1 promote hematopoietic progenitor homing to bone marrow. Proc Natl Acad Sci U S A. 1998. doi:10.1073/pnas.95.24.14423

61. Cancilla D, Thakellapalli H, Meyers MJ, et al. Targeting CXCR4, VLA4, and CXCR2 for Hematopoietic Stem Cell Mobilization. Blood. 2019;134(Supplement_1). doi:10.1182/blood-2019-124373

62. Keller JR, Ortiz M, Ruscetti FW. Steel factor (c-kit ligand) promotes the survival of hematopoietic stem/progenitor cells in the absence of cell division. Blood. 1995. doi:10.1182/blood.v86.5.1757.bloodjournal8651757

63. Kenswil KJG, Jaramillo AC, Ping Z, et al. Characterization of Endothelial Cells Associated with Hematopoietic Niche Formation in Humans Identifies IL-33 As an Anabolic Factor. Cell Rep. 2018;22(3). doi:10.1016/j.celrep.2017.12.070

64. Christodoulou C, Spencer JA, Yeh SCA, et al. Live-animal imaging of native haematopoietic stem and progenitor cells. Nature. 2020. doi:10.1038/s41586-020-1971-z

65. Ema H, Morita Y, Suda T. Heterogeneity and hierarchy of hematopoietic stem cells. Exp Hematol. 2014;42(2). doi:10.1016/j.exphem.2013.11.004

66. Gauntner TD, Brunstein CG, Cao Q, et al. Association of CD34 Cell Dose with 5-Year Overall Survival after Peripheral Blood Allogeneic Hematopoietic Cell Transplantation in Adults with Hematologic Malignancies. Transplant Cell Ther. 2022;28(2). doi:10.1016/j.jtct.2021.11.004

67. Jakubowski AA, Small TN, Kernan NA, et al. T cell-depleted unrelated donor stem cell transplantation provides favorable disease-free survival for adults with hematologic malignancies. Biol Blood Marrow Transplant. 2011;17(9). doi:10.1016/j.bbmt.2011.01.005

68. Oubari F, Amirizade N, Mohammadpour H, Nakhlestani M, Zarif MN. The important role of FLT3-L in ex vivo expansion of hematopoietic stem cells following co-culture with mesenchymal stem cells. Cell J. 2015;17(2). doi:10.22074/cellj.2016.3715

69. Serke S, Watts M, Knudsen LM, et al. In-vitro clonogenicity of mobilized peripheral blood CD34-expressing cells: Inverse correlation to both relative and absolute numbers of CD34-expressing cells. Br J Haematol. 1996;95(2). doi:10.1046/j.1365-2141.1996.d01-1918.x

70. Berglund S, Magalhaes I, Gaballa A, Vanherberghen B, Uhlin M. Advances in umbilical cord blood cell therapy: the present and the future. Expert Opin Biol Ther. 2017. doi:10.1080/14712598.2017.1316713

71. De la Morena MT, Gatti RA. A History of Bone Marrow Transplantation. Hematol Oncol Clin North Am. 2011;25(1). doi:10.1016/j.hoc.2010.11.001

72. D’Souza A, Fretham C, Lee SJ, et al. Current Use of and Trends in Hematopoietic Cell Transplantation in the United States. Biol Blood Marrow Transplant. 2020;26(8). doi:10.1016/j.bbmt.2020.04.013

73. Delaney C, Heimfeld S, Brashem-Stein C, Voorhies H, Manger RL, Bernstein ID. Notch-mediated expansion of human cord blood progenitor cells capable of rapid myeloid reconstitution. Nat Med. 2010. doi:10.1038/nm.2080

74. Fares I, Chagraoui J, Gareau Y, et al. Pyrimidoindole derivatives are agonists of human hematopoietic stem cell self-renewal. Science (80-). 2014. doi:10.1126/science.1256337

75. Milano F, Heimfeld S, Riffkin IB, et al. Infusion of a Non HLA-Matched Off-the-Shelf Ex Vivo Expanded Cord Blood Progenitor Cell Product Following Myeloablative Cord Blood Transplantation Is Safe, Decreases the Time to Hematopoietic Recovery, and Results in Excellent Overall Survival. Blood. 2014. doi:10.1182/blood.v124.21.46.46

76. Horwitz ME, Wease S, Blackwell B, et al. Phase I/II study of stem-cell transplantation using a single cord blood unit expanded ex vivo with nicotinamide. J Clin Oncol. 2019. doi:10.1200/JCO.18.00053

77. de Lima M, McMannis J, Gee A, et al. Transplantation of ex vivo expanded cord blood cells using the copper chelator tetraethylenepentamine: A phase I/II clinical trial. Bone Marrow Transplant. 2008. doi:10.1038/sj.bmt.1705979

78. Wagner JE, Brunstein CG, Boitano AE, et al. Phase I/II Trial of StemRegenin-1 Expanded Umbilical Cord Blood Hematopoietic Stem Cells Supports Testing as a Stand-Alone Graft. Cell Stem Cell. 2016. doi:10.1016/j.stem.2015.10.004

79. Choi JS, Mahadik BP, Harley BAC. Engineering the hematopoietic stem cell niche: Frontiers in biomaterial science. Biotechnol J. 2015. doi:10.1002/biot.201400758

80. Liu B, Jin M, Wang DA. In vitro expansion of hematopoietic stem cells in a porous hydrogel-based 3D culture system. Acta Biomater. 2023. doi:10.1016/j.actbio.2023.01.057

81. Kowalczyk M, Waldron K, Kresnowati P, Danquah MK. Process challenges relating to hematopoietic stem cell cultivation in bioreactors. J Ind Microbiol Biotechnol. 2011. doi:10.1007/s10295-011-0951-6

82. Rodrigues CAV, Fernandes TG, Diogo MM, da Silva CL, Cabral JMS. Stem cell cultivation in bioreactors. Biotechnol Adv. 2011;29(6). doi:10.1016/j.biotechadv.2011.06.009

83. Prielhofer R, Lindner C, Sohail A, Reininger M, Graumann K. Development of Scalable Stem Cell Cultivation Processes in Bioreactors. Chemie Ing Tech. 2022;94(9). doi:10.1002/cite.202255185

84. Meissner P, Schröder B, Herfurth C, Biselli M. Development of a fixed bed bioreactor for the expansion of human hematopoietic progenitor cells. Cytotechnology. 1999;30(1-3). doi:10.1023/a:1008085932764

85. Chaudhuri O, Gu L, Klumpers D, et al. Hydrogels with tunable stress relaxation regulate stem cell fate and activity. Nat Mater. 2016;15(3). doi:10.1038/nmat4489

86. Trappmann B, Gautrot JE, Connelly JT, et al. Extracellular-matrix tethering regulates stem-cell fate. Nat Mater. 2012. doi:10.1038/nmat3339

87. Khetan S, Guvendiren M, Legant WR, Cohen DM, Chen CS, Burdick JA. Degradation-mediated cellular traction directs stem cell fate in covalently crosslinked three-dimensional hydrogels. Nat Mater. 2013;12(5). doi:10.1038/nmat3586

88. Baker BM, Trappmann B, Wang WY, et al. Cell-mediated fibre recruitment drives extracellular matrix mechanosensing in engineered fibrillar microenvironments. Nat Mater. 2015. doi:10.1038/nmat4444

89. You Y, Kobayashi K, Colak B, et al. Engineered cell-degradable poly(2-alkyl-2-oxazoline) hydrogel for epicardial placement of mesenchymal stem cells for myocardial repair. Biomaterials. 2021;269. doi:10.1016/j.biomaterials.2020.120356

90. Rao V V., Wechsler ME, Cravens E, et al. Granular PEG hydrogels mediate osteoporotic MSC clustering via N-cadherin influencing the pro-resorptive bias of their secretory profile. Acta Biomater. 2022;145. doi:10.1016/j.actbio.2022.04.023

91. De Falco E, Porcelli D, Torella AR, et al. SDF-1 involvement in endothelial phenotype and ischemia-induced recruitment of bone marrow progenitor cells. Blood. 2004;104(12). doi:10.1182/blood-2003-12-4423

92. Santiago B, Calonge E, Rey MJD, et al. CXCL12 gene expression is upregulated by hypoxia and growth arrest but not by inflammatory cytokines in rheumatoid synovial fibroblasts. Cytokine. 2011;53(2). doi:10.1016/j.cyto.2010.06.006

93. Ceradini DJ, Kulkarni AR, Callaghan MJ, et al. Progenitor cell trafficking is regulated by hypoxic gradients through HIF-1 induction of SDF-1. Nat Med. 2004;10(8). doi:10.1038/nm1075

94. Flores-Guzmán P, Gutiérrez-Rodríguez M, Mayani H. In vitro proliferation, expansion, and differentiation of a CD34+ cell-enriched hematopoietic cell population from human umbilical cord blood in response to recombinant cytokines. Arch Med Res. 2002. doi:10.1016/S0188-4409(01)00368-X

95. Petaroudi M, Rodrigo-Navarro A, Dobre O, Dalby MJ, Salmeron-Sanchez M. Living Biomaterials to Engineer Hematopoietic Stem Cell Niches. Adv Healthc Mater. 2022;11(20). doi:10.1002/adhm.202200964

96. Takubo K, Nagamatsu G, Kobayashi CI, et al. Regulation of glycolysis by Pdk functions as a metabolic checkpoint for cell cycle quiescence in hematopoietic stem cells. Cell Stem Cell. 2013;12(1). doi:10.1016/j.stem.2012.10.011

97. Spencer JA, Ferraro F, Roussakis E, et al. Direct measurement of local oxygen concentration in the bone marrow of live animals. Nature. 2014;508(7495):269–273. doi:10.1038/nature13034

98. Preciado S, Sirerol-Piquer M aS, Muntión S, et al. Co-administration of human MSC overexpressing HIF-1α increases human CD34+ cell engraftment in vivo. Stem Cell Res Ther. 2021;12(1). doi:10.1186/s13287-021-02669-z

99. Peng L, Nadal C, Gautrot JE. Growth of mesenchymal stem cells at the surface of silicone, mineral and plant-based oils. Biomed Mater. 2023;18(3). doi:10.1088/1748-605x/acbdda

100. Pfaffl MW. A new mathematical model for relative quantification in real-time RT–PCR. Nucleic Acids Res. 2001;29(9). doi:10.1093/nar/29.9.e45

