## Supplementary Information for "Biomimetic Artificial Bone Marrow Niches for the Scale Up of Hematopoietic Stem Cells"

**Supplementary Videos S1 and S2.** MSCs and HSPCs interact in co-culture. Time-lapse videos showing unstained MSCs growing on 2D (Video S1) or bioemulsions (Video S2) for 7 days and freshly seeded HSCs stained with cell tracker (green). Images were taken every 5 minutes for a total period of 9 hours.

**Supplementary Video S3.** 3D sectioning through two-photon microscopy of @BMN with HSPCs cultured for 15 days. Bioemulsions were formed with fluorescently tagged PLL (red) to visualise the interface and cells were stained to visualise the nuclei (blue) and the actin cytoskeleton (green).

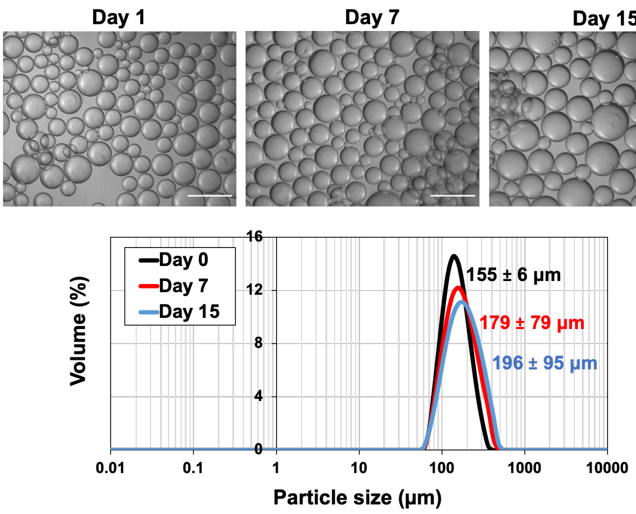


**Supplementary Figure S1.** PLL-stabilised bioemulsions remain stable under agitation for 15 days. The diameter of fluorinated oil bioemulsions was evaluated at different time points (Mastersizer), under agitation on an orbital shaker at 60 rpm and 37ºC. Top, brightfield images illustrate representative distributions of droplet sizes. Scale bar, 200 µm. Bottome, particle size distribution at days 0, 7 and 15. The extrapolated diameters are indicated as means ± standard deviations. N = 3.

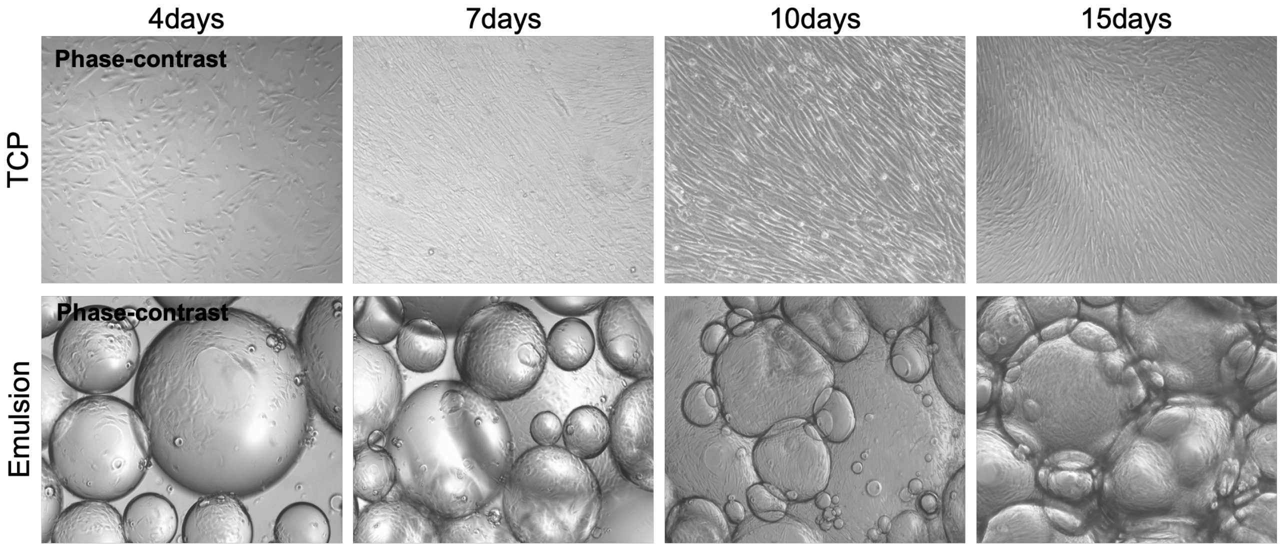


**Supplementary Figure S2.** Proliferation of MSCs at the surface of TCP and bioemulsions over the course of 15 days in MSC expansion medium (phase contrast images).

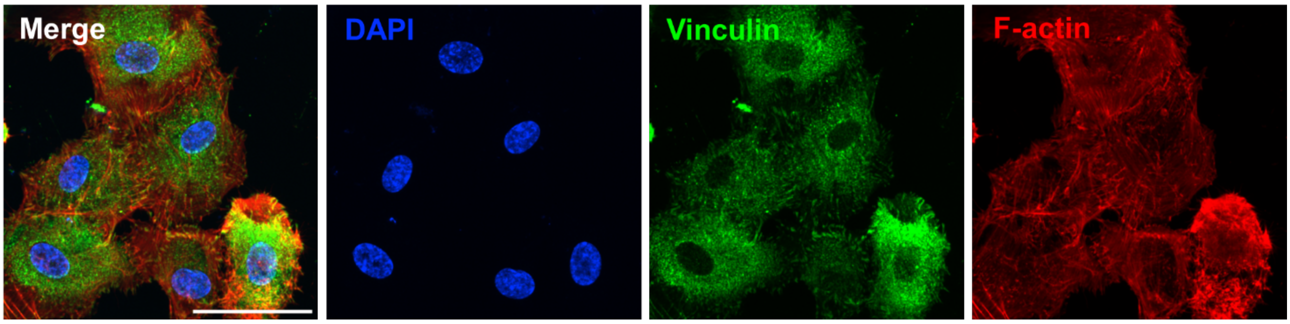


**Supplementary Figure S3**. MSCs were cultured on glass substrates 3 days prior to fixation and immunostaining. Cells were stained with vinculin (green) and Phalloidin (red) Scale bar, 20 µm.

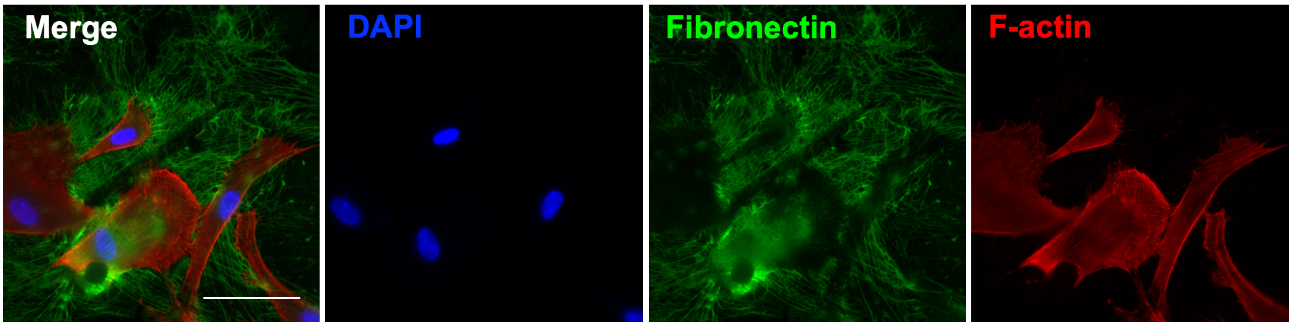


**Supplementary Figure S4.** MSCs adhering to fibronectin coated PLL nanosheets further remodel and assemble fibronectin into a fibrillar matrix. MSCs were cultured on bioemulsions 5 days prior to fixation and immunostaining. Note that cells have rearranged and/or secreted fibronectin at interfaces prior to migrating to other areas, contributing to the overall remodelling of the niche, rather than simply secreting pericellular matrix. Scale bar, 20 µm.

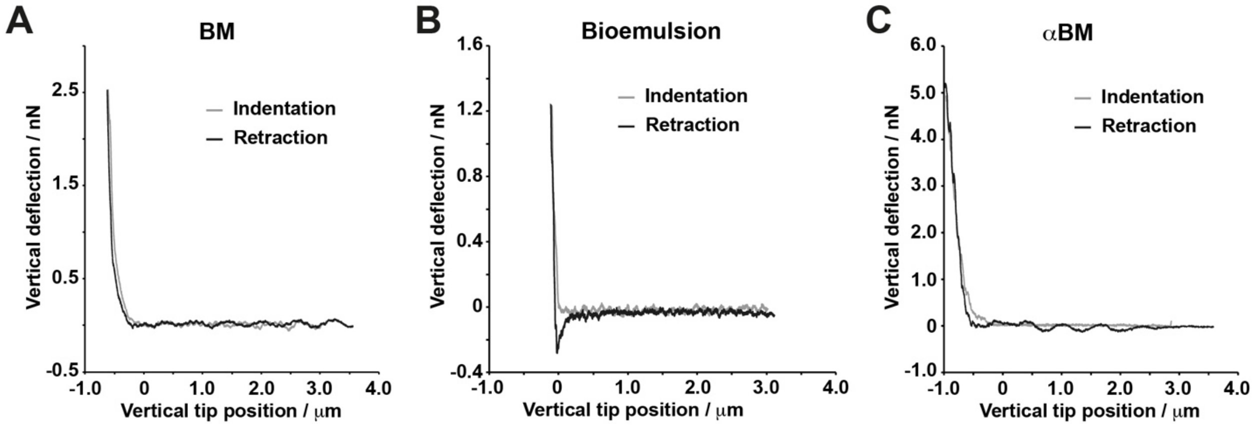


**Supplementary Figure S5.** Representative examples of AFM indentation and retraction traces obtained for bone marrow samples (BM), PLL nanosheet stabilised microdroplets (bioemulsions) and artificial bone marrow samples (@BM).

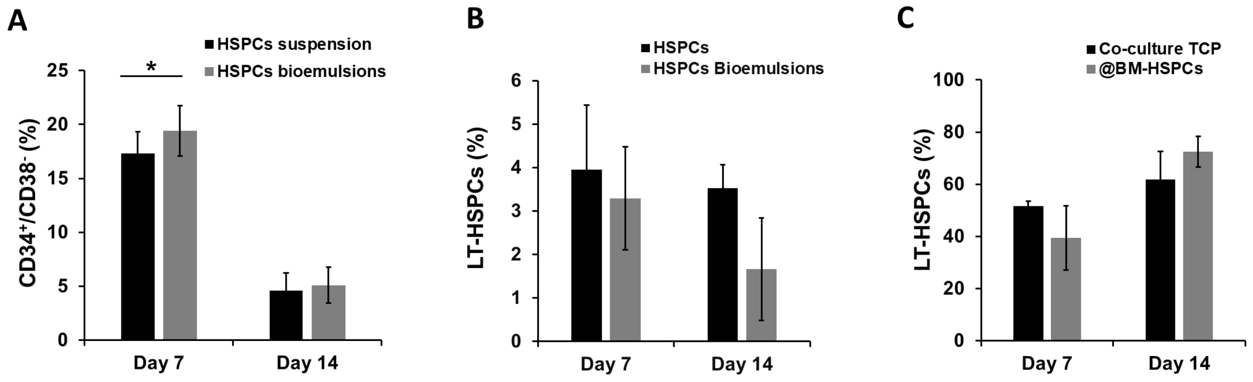


**Supplementary Figure S6.** (A) Quantification of the densities of $CD34^+/CD38^-$ cells determined by flow cytometry after mono-culture of HSPCs for 14 days in suspension or on bioemulsions. Mean ± SD; N = 3; (*, P,0.05). After doublet exclusion, mononuclear cells were gated based on low/high expression of calcein to identify live cells and exclude bioemulsions and dead cells. CD34 and CD38 markers were used to identify the population of interest. (B) Percentage of LT-HSCs quantified by flow cytometry after culturing HSPCs for 14 days in suspension or on bioemulsions without MSCs. Mean ± SD; N = 3; (no statistically significant differences observed). After doublet exclusion, mononuclear cells were gated based on low/high expression of calcein to identify live cells and exclude bioemulsions and dead cells. $CD34^+/CD38^-$ subset was selected and $CD90^+/CD45RA^-$ cells were subsequently gated to identify the population of interest. (C) Percentage of LT-HSCs quantified by flow cytometry after culturing HSPCs for 14 days in co-culture either on TCP or on bioemulsions. Mean ± SD; N = 3; no statistically significant differences observed. After doublet exclusion, mononuclear cells were gated based on low/high expression of calcein to identify live cells and exclude bioemulsions and dead cells. $CD34^+/CD38^-$ subset was selected and $CD90^+/CD45RA^-$ cells were subsequently gated to identify the population of interest.

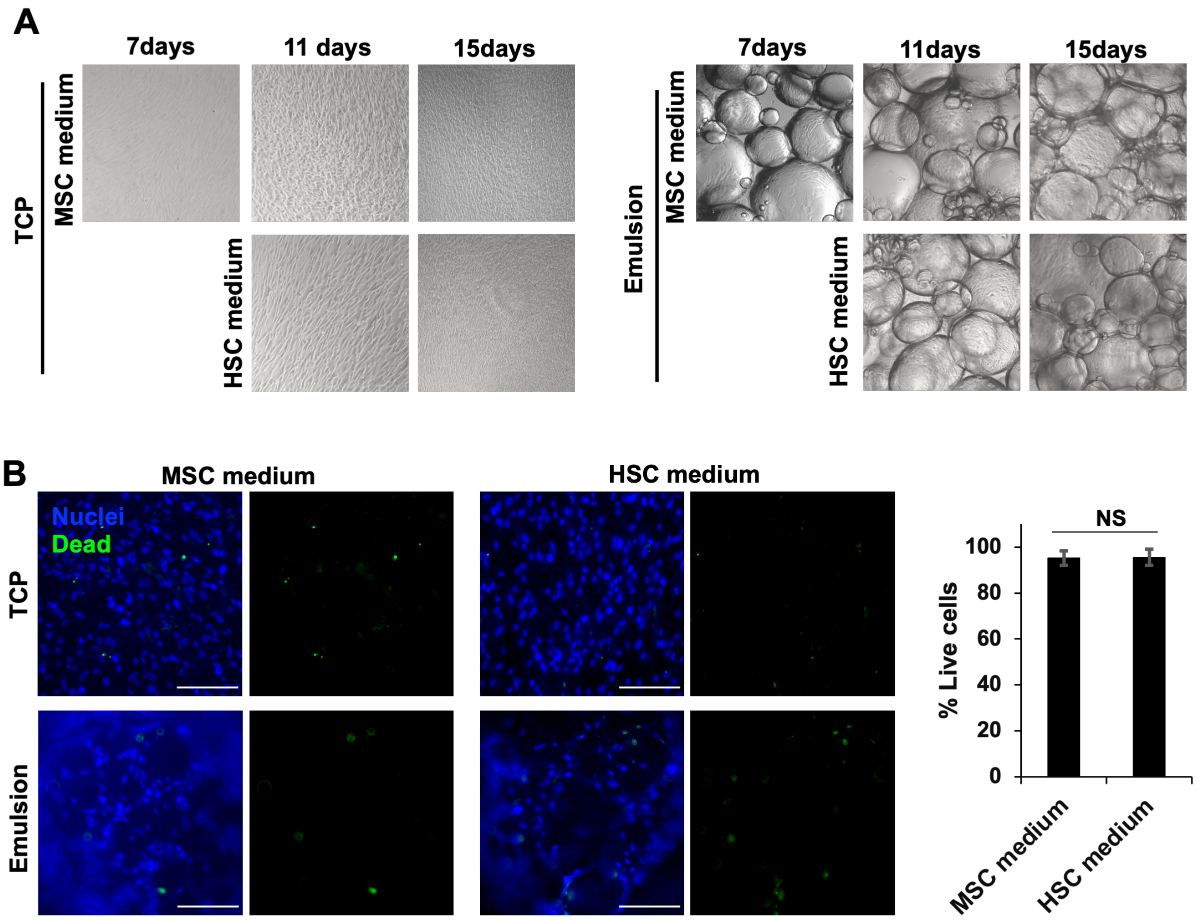


**Supplementary Figure S7.** HSC medium does not influence the viability of MSCs. A) Brightfield images of MSCs cultured in MSC and HSC media on TCP and bioemulsions. MSC medium was switched (or not) with HSC medium at day 7. B) Left, MSCs growing on 2D TCP and bioemulsions for 15 days in MSC or HSC medium, stained with EthD1 (green) to visualise dead cells. Scale bar, 200 µm. Right, quantification of percentages of live MSCs on bioemulsions cultured in MSC and HSC media. Mean ± SD; N = 3; NS, P > 0.05.

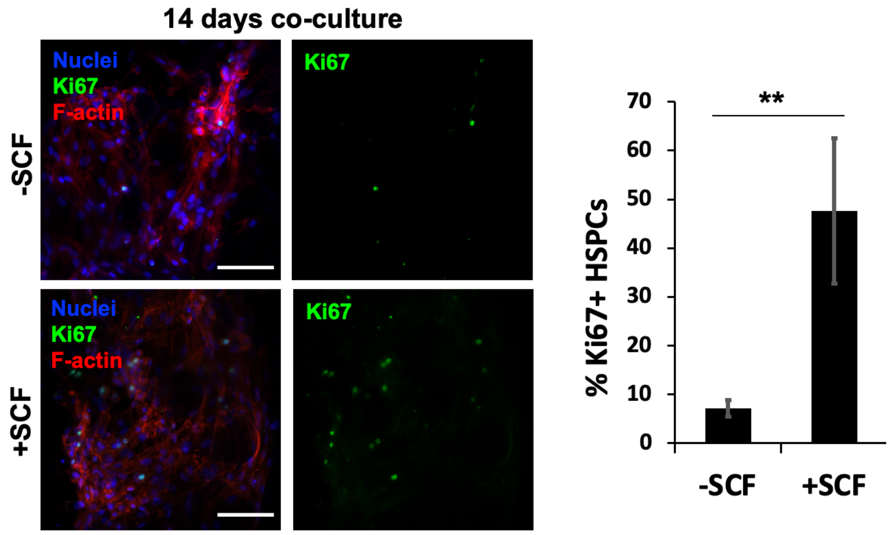


**Supplementary Figure S8.** SCF induces the proliferation of HSPCs. Left, z-Projections of confocal z-stacks of co-cultures growing on bioemulsions, immunostained for Ki-67 to identify actively cycling cells. Scale bar, 50 µm. Right, quantification of percentage of Ki-67$^{+}$ HSPCs on bioemulsions after 14 days of co-culture in HSC medium (with and without SCF). Mean ± SD; N = 3; **, $P < 0.01$.

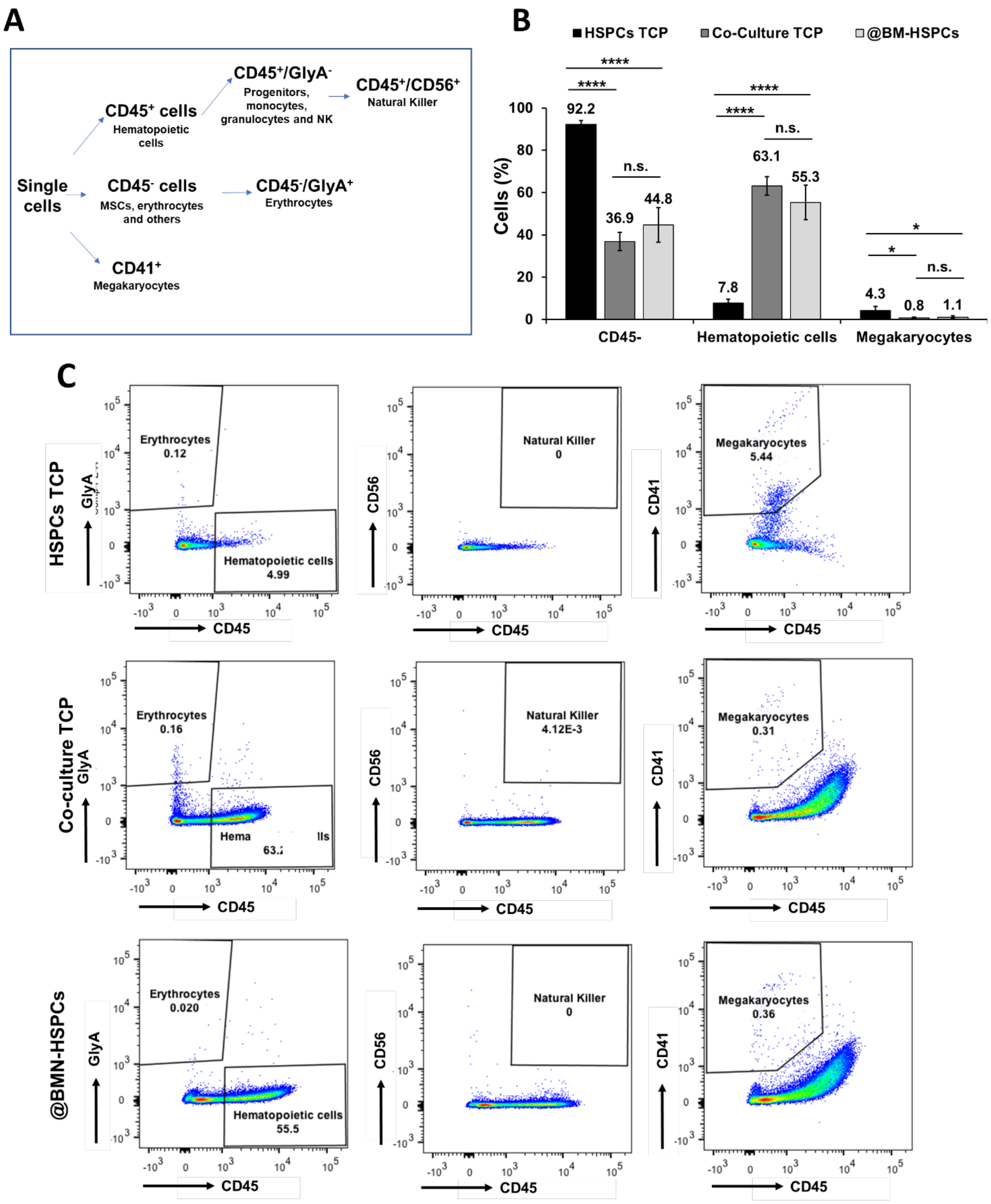


**Supplementary Figure S9.** (A) Schematics of the gating strategy to identify hematopoietic subpopulations after 14 days of culture. (B) Percentage of each subpopulation obtained by flow cytometry after culturing HSPCs for 14 days in suspension, or in co-culture on TCP or bioemulsion. Mean ± SD; N = 4; (*, P,0.05; **, P < 0.01). (C) Gating strategy for identification of cell populations of interest in HSCPs grown for 14 days in suspension (upper panels), in co-culture on TCP (mid panels), or on bioemulsion (lower panels). After doublet exclusion, mononuclear cells were gated based on low/high CD45 expression to identify hematopoietic cells. Erythrocytes were identified as $CD45^-/Gly^+$, NK were identified as $CD45^+/CD56^-$, and Megakaryocytes were identified as $CD45^-/CD41^+$.

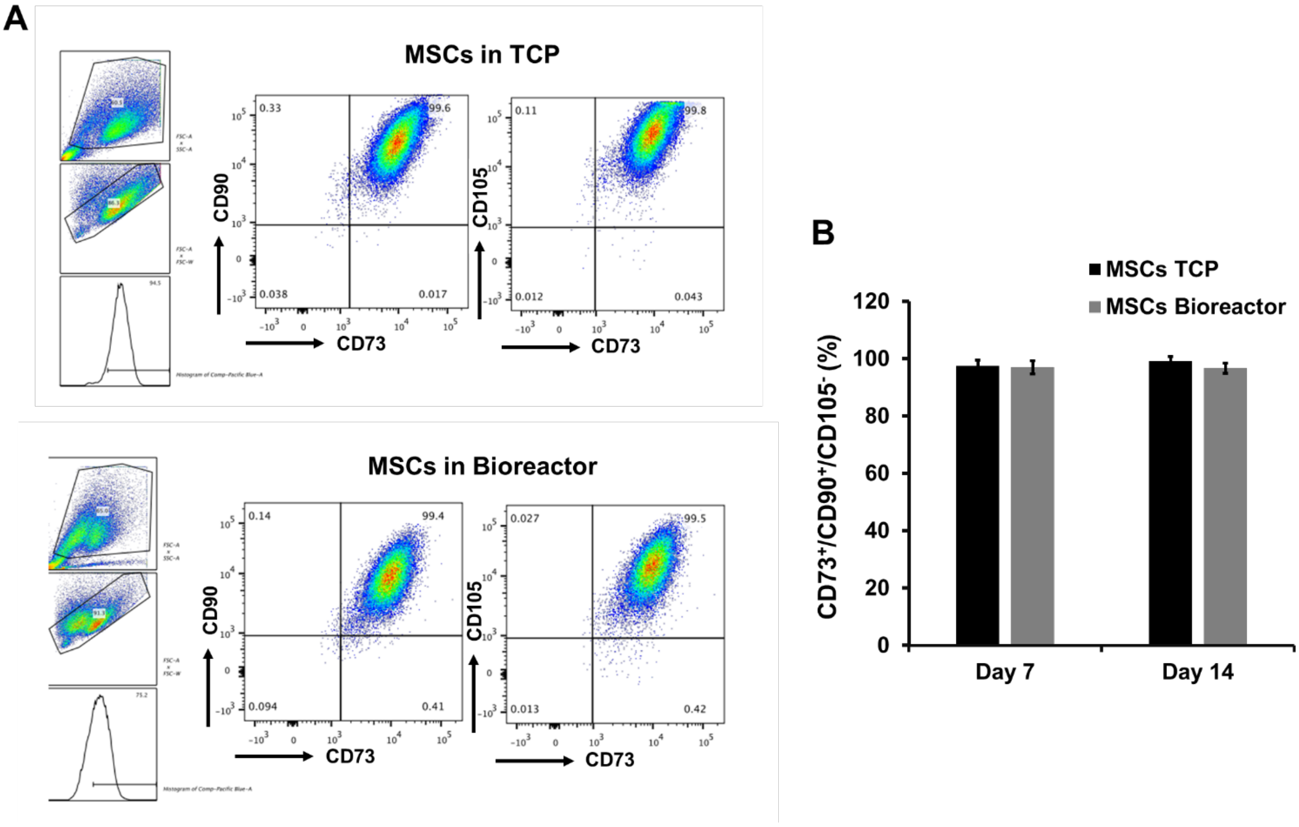


**Supplementary Figure S10.** (A) Gating strategy to identify MSCs expressing key markers after 14 days of culture growing both on TCP (upper panels) and on bioemulsions in conical flask bioreactor format (lower panels). After doublet exclusion, mononuclear cells were gated based on low/high expression of calcein to identify live cells and exclude bioemulsions and dead cells. $CD73^+/CD90^+$ subset was selected and $CD105^+$ cells were subsequently gated to identify the population of interest. (B) Graph show percentage of $CD73^+/CD90^+/$ $CD105^+$ obtained by flow cytometry after culturing MSCs for 7 and 14 days on TCP or on bioemulsions in a bioreactor. Mean ± SD; N = 3; (no statistically significant differences observed).

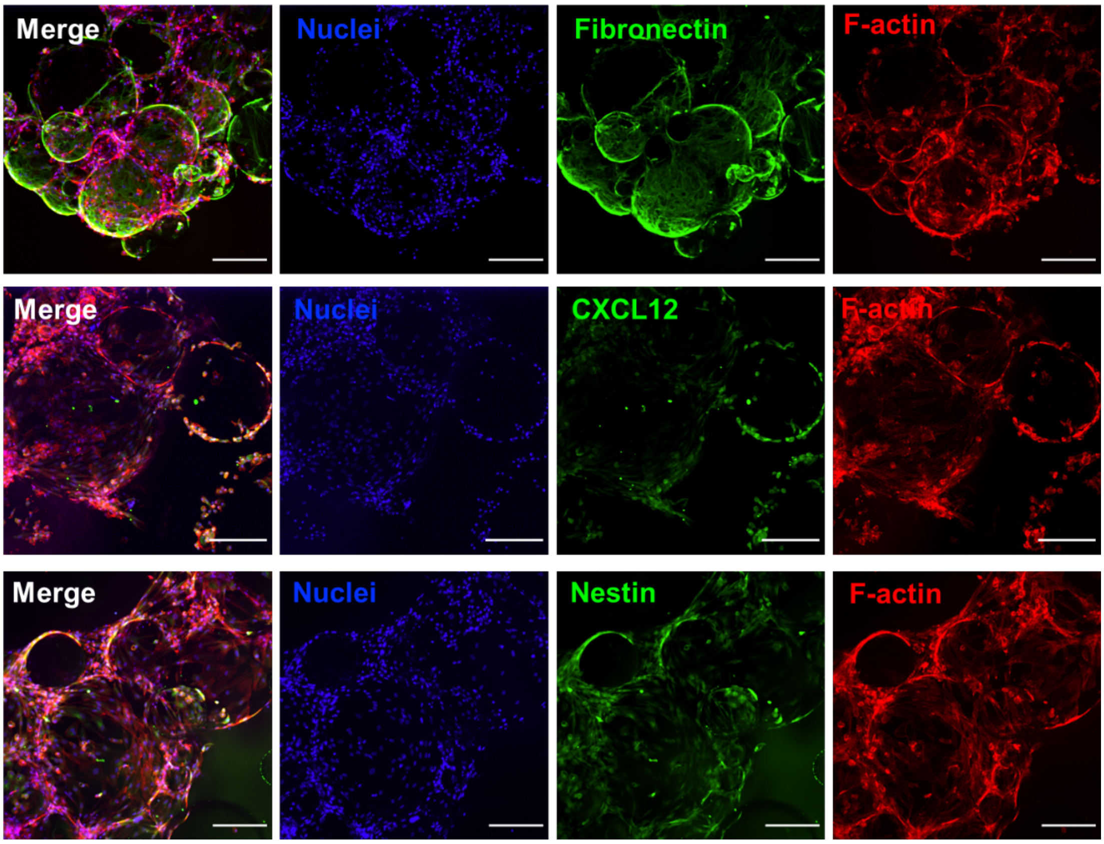


**Supplementary Figure S11.** z-Projection of confocal z-stack of co-cultured @BMN grown in conical flask bioreactor for 14 days. Scale bar, 100 µm.

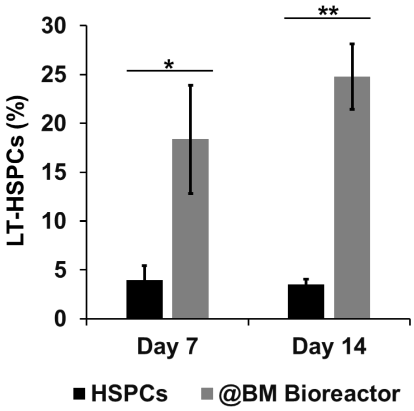


**Supplementary Figure S12.** Percentage of LT-HSCs obtained by flow cytometry after culturing HSPCs for 14 days in suspension or within @BMNs in conical flask bioreactors. Mean ± SD; N = 3; (*, P,0.05: **, P<0.001). After doublet exclusion, mononuclear cells were gated based on low/high expression of calcein to identify live cells and exclude bioemulsions and dead cells. $CD34^+/CD38^-$ subset was selected and $CD90^+/CD45RA^-$ cells were subsequently gated to identify the population of interest.

**Details of Taqman probes used for target genes for qPCR.**

| Gene | Assay |
|---|---|
| **VCAM-1** | Hs01003372_m1 |
| **Interleukin -6** | Hs00174131_m1 |
| **Thrombopoietin** | Hs01061346_m1 |
| **Angiopoietin 1** | Hs00181613_m1 |
| **CXCL2** | Hs03676656_mH |
| **Stem Cell Factor** | Hs01070032_m1 |
| **Jagged 1** | Hs00241497_m1 |
| **Nestin** | Hs04187831_g1 |
